# Emergence of hyper insecticide-resistant dengue vectors in Indochina Peninsula: threats of concomitant knockdown resistance mutations

**DOI:** 10.1101/2022.03.05.483084

**Authors:** Shinji Kasai, Kentaro Itokawa, Nozomi Uemura, Aki Takaoka, Shogo Furutani, Yoshihide Maekawa, Daisuke Kobayashi, Nozomi Imanishi-Kobayashi, Michael Amoa-Bosompem, Katsunori Murota, Yukiko Higa, Hitoshi Kawada, Noboru Minakawa, Tran Chi Cuong, Nguyen Thi Yen, Tran Vu Phong, Sath Keo, Kroesna Kang, Kozue Miura, Lee Ching Ng, Hwa-Jen Teng, Samuel Dadzie, Sri Subekti, Kris Cahyo Mulyatno, Kyoko Sawabe, Takashi Tomita, Osamu Komagata

**Author notes:** These authors contributed equally (Shinji Kasai, Kentaro Itokawa, Nozomi Uemura). Corresponding authors: Name: Shinji Kasai, Ph.D., Address: 1-23-1 Toyama, Shinjukuku, Tokyo 162–8640, Japan, Name: Osamu Komagata, Ph.D., Address: 1-23-1 Toyama, Shinjukuku, Tokyo 162–8640, Japan. **Author Contributions:** S. Kasai, O.K., K.I., T.V.P., K.S., and T.T. designed the research; S. Kasai, N.U., K.I., S.F., A.T., and H.K. performed research; S. Kasai, K.I., Y.M., D.K., M.A.B., N.I.K., K.M., Y.H., H.K., N.M., T.C.C., N.T.Y., T.V.P., H.J.T., L.C.N, S.D., K.K., S. Keof, K.M, S.S.B., K.C.M, and K.S. contributed new reagents/analytic tools; S. Kasai, O.K., H.K., and K.I. analyzed the data; S. Kasai, O.K., K.I., N.U., A.T., S.F., Y.M., D.K., M.A.B., N.I.K., K.M., Y.H., H.K., N.M., T.C.C., N.T.Y., T.V.P., H.J.T., S.D., K.K., S. Keof, K.M., K.S., S.S.B., K.C.M, T.T., and L.C.N. wrote the manuscript.

## Abstract

*Aedes aegypti* (Linnaeus, 1762) is the main mosquito vector for dengue and other arboviral infectious diseases. Control of this important vector highly relies on the use of insecticides, especially pyrethroids. Nevertheless, the development of pyrethroid resistance is a major obstacle to mosquito/disease control worldwide. Here, we focused on the mutations in the target site of pyrethroid insecticides, voltage-sensitive sodium channel (*Vssc*), and found that *Ae. aegypti* collected from Vietnam has the L982W allele in the *Vssc* at a high frequency (>79%). L982W mutation is located in the highly conserved region of *Vssc* that is associated with sodium–ion selectivity and permeation rate. Strains having the L982W allele showed similar or even higher levels of resistance to pyrethroids than those having V1016G, a typical knockdown resistance allele in Asia. Furthermore, concomitant mutations L982W+F1534C and V1016G+F1534C were confirmed, and strains having these multiple *Vssc* mutations exhibited incomparably higher levels of pyrethroid resistance than any other field population ever reported. Molecular modeling analysis confirmed that these concomitant mutant alleles could interfere with approaching pyrethroid to *Vssc*. Remarkably, >90% of *Vssc* of *Ae. aegypti* were occupied by these hyper insecticide-resistant haplotypes in Phnom Penh city, Cambodia. Analysis of whole *Vssc* coding genes suggested that *Vssc*s have evolved into stronger resistant forms efficiently through gene recombination events. At this point, L982W has never been detected in *Vssc* of *Ae. aegypti* from any other neighboring countries. We strongly emphasize the need to be vigilant about these strong resistance genes spreading to the world through Indochina Peninsula.

**Significance Statement:** The high frequency (>78%) of the L982W allele was detected at the target site of the pyrethroid insecticide, the voltage-sensitive sodium channel (*Vssc*) of *Aedes aegypti* collected from Vietnam and Cambodia. Haplotypes having concomitant mutations L982W+F1534C and V1016G+F1534C were also confirmed in both countries, and their frequency was high (>90%) in Phnom Penh, Cambodia. Strains having these haplotypes exhibited substantially higher levels of pyrethroid resistance than any other field population ever reported. The L982W mutation has never been detected in any country of the Indochina Peninsula except Vietnam and Cambodia, but it may be spreading to other areas of Asia, which can cause an unprecedentedly serious threat to the control of dengue fever as well as other *Aedes*-borne infectious diseases.

## Introduction

*Aedes aegypti* is a vector for a variety of arboviral infectious diseases including dengue, chikungunya, zika, and yellow fever. Dengue is the most prevalent among them; the number of dengue cases has increased thirty-fold in the past 50 years (1). One of the factors for this is contributed by the geographical expansion of *Ae. aegypti* due to climate change (2). A modeling study estimated that 390 million people are infected annually (3). Thus, the WHO has designated dengue as one of the top 10 threats to global health in 2019, together with other important issues such as antimicrobial resistance and global influenza pandemic (https://www.who.int/news-room/spotlight/ten-threats-to-global-health-in-2019). Various new technologies, including the use of sterile insect technique and transgenic- and Wolbachia infected-mosquitoes, have been developed to break the transmission of dengue virus (4–9). However, they are not ready for large-scale implementation, and there are also some arguments on the release of genetically engineered organisms to the environment (10). With no cost-effective vaccine or medication available, the use of vector control insecticides remains the only approach to manage dengue epidemics. Conversely, many *Ae. aegypti* populations have developed resistance to pyrethroids, making vector control more difficult (11).

One of the major mechanisms of pyrethroid resistance is the reduced sensitivity of the target site, i.e., voltage-sensitive sodium channel (*Vssc*). *Vssc* gene mutations that cause amino acid substitutions and confer pyrethroid resistance are collectively called knockdown resistance or *kdr* (12-16). Several *kdr* mutations (alleles) including V253F, V410L, I1011M, V1016G, and F1534C have been reported in *Ae. aegypti* (17-21). In Asia, V1016G and F1534C are recognized as major *kdr* alleles. Co-occurrences of alleles S989P, V1016G, and F1534C are also reported in *Ae. aegypti* collected from Thailand, Myanmar, Indonesia, China, Sri Lanka, Saudi Arabia, Lao, and Malaysia (22-27). Although electrophysiological studies revealed that triple mutations of S989P, V1016G, and F1534C strongly reduced the susceptibility of mosquitoes to permethrin and deltamethrin (28), its phenotypic effect on the insecticide resistance of insects has not been elucidated at all.

In Cambodia, there were two major dengue endemics in 2007 and 2012 with 39,618 and 42,362 reported cases, respectively. Nearly 600 deaths were also recorded within these years (29, 30). Dichlorodiphenyltrichloroethane (DDT) had been used to control dengue and malaria in this country followed by permethrin and deltamethrin treatments by thermal fogging and ultra-low volume (ULV) spraying (31). It is also reported that *Ae. aegypti* collected in and around Phnom Penh city exhibited an extremely high level of resistance to several pyrethroids. The mortality rates of *Ae. aegypti* population to permethrin and deltamethrin in the Phnom Penh were 0 and less than 10%, respectively, by WHO diagnostic method (31). In 2022, Boyer et al. also reported similar results using seven pyrethroid insecticides (32). Nevertheless, the molecular mechanism of pyrethroid resistance of *Ae. aegypti* collected from Cambodia remains unknown. It is crucial to monitor the insecticide resistance in *Aedes aegypti* collected from dengue epidemic area and to elucidate molecular mechanisms of insecticide resistance of this important mosquito species.

In this study, we examined the relationship between *Vssc* and pyrethroid resistance in *Ae. aegypti* collected in Asia. Ten new isogenic *kdr* strains of *Ae. aegypti* were established and the susceptibility of each strain to pyrethroids was examined. Full-length *Vssc* coding DNA sequences were determined for all strains, which resulted in the identification of several new amino acid substitutions. We focused on the co-occurrence of *kdr* mutations that confer an extremely high level of pyrethroid resistance. Furthermore, molecular evolution processes of *Vssc* to accumulate *kdr* alleles were discussed.

## Results

### Permethrin susceptibility

Permethrin susceptibility of 23 *Ae. aegypti* populations collected from Vietnam, Indonesia, Ghana, and Taiwan as well as an insecticide-susceptible strain (SMK) was evaluated (Fig. 1*A*). This study was conducted as a part of a project to assess pyrethroid susceptibility of world *Aedes* mosquitoes (33). Diagnostic doses were established on the basis of the susceptibility of the reference *Aedes albopictus* (Skuse, 1895) strain HKM (33). The mortality of the SMK strain at 5.9 ng permethrin per female, which is equivalent to 99 percent lethal dose (LD_99_) of SMK, was 100% (Fig. 1*A*), suggesting that this dose of permethrin was also effective in evaluating the resistance status of *Ae. aegypti*. By contrast, the field populations showed lower mortalities across the board. When exposed to 5.9 ng, all populations except for three Taiwanese and a Ghana population (Aburi) had less than 20% mortality. Raising the dose by 10 times (59 ng) led to higher mortality for all strains. Nonetheless, only two populations from Ghana had 100% mortality. Even the Taiwanese populations were found to have surviving individuals. Particularly, the populations of Hanoi 2 (Vietnam) and Simo Sidomulyo (Indonesia) showed mortality below 30% even when exposed to 59 ng permethrin.

**Figure 1.**
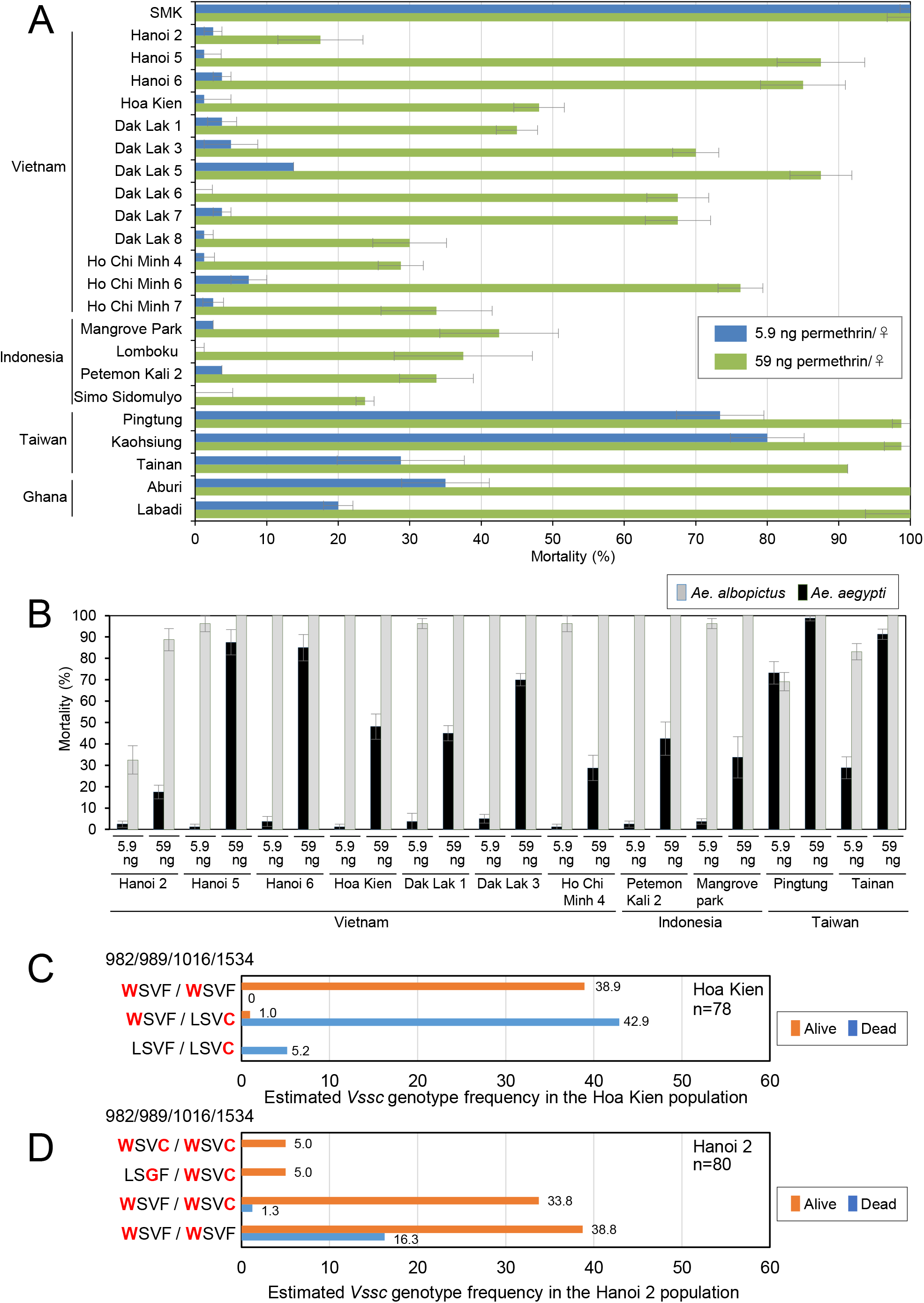
Permethrin susceptibility and frequency of *kdr* alleles in adult *Aedes aegypti* collected from four countries. (*A*) Mortalities of adult *Ae. aegypti* after exposures to permethrin. Mosquitoes were topically exposed to two doses of permethrin, i.e., 5.9 or 59 ng, which were respectively equivalent to the LD_99_ and LD_99_×10 doses for the susceptible *Ae. albopictus* strain HKM (33). Twenty females were treated in each assay, whereby the subsequent proportion of dead mosquitoes was calculated 24 h after the treatment. Data are the average of four replicates. Error bars indicate standard errors for the mortalities of each population. (*B*) Comparison of permethrin susceptibility between *Ae. aegypti* and *Ae. albopictus* which were collected from the same areas. Mortality data for *Ae. albopictus* was from the previous report (33). (*C*, *D*) Genotyping of *Vssc* genes from dead and surviving mosquitoes after the bioassay with 59 ng permethrin. The total number of mosquitoes tested from Hoa Kien (*C*) and Hanoi 2 (*D*) were 78 and 80, respectively.

### *Ae. aegypti* develops higher levels of resistance to permethrin than *Ae. albopictus*

We have previously tested the permethrin susceptibility of *Ae. albopictus*, another major vector mosquito for dengue, chikungunya, and zika, collected from four countries between 2015 and 2017 (33), and the permethrin susceptibilities of these two *Aedes* species collected from the same larval sources were compared (Fig. 1*B*). Except for the Taiwanese Pingtung population, all *Ae. aegypti* populations showed lower mortalities than *Ae. albopictus*.

### Associations between *Vssc* alleles and permethrin susceptibility

To evaluate the association between *Vssc* alleles and pyrethroid susceptibility, genotyping analysis was conducted on individual mosquitoes of the two Vietnamese *Ae. aegypti* populations (Hoa Kien and Hanoi 2) that were subjected to 59 ng permethrin treatment. We first attempted to genotype 3 loci containing well-known variants, namely, S989P, V1016G, and F1534C, which have been shown to confer pyrethroid resistance (11, 17, 28), especially among Asian *Ae. aegypti*. In this process, we found that an additional known variant L982W was included in the same polymerase chain reaction (PCR) product for S989P and V1016G (Fig. 1*C* and 1*D*). V1016G allele, one of the major *kdr* variants, was not detected from these two Vietnamese populations. Contrarily and unexpectedly, the Hoa Kien population had L982W at a high frequency of 72.7%. There was a strong genotype–phenotype correlation, where 97.6% (40/41) of the survived individuals had L982W homozygously, whereas 89.2% (33/37) of the dead individuals had L982W and F1534C both heterozygously (Fig. 1*C*). Although the frequency of F1534C in the Hoa Kien population was 27.3%, no individuals homozygous for L982W had an F1534C allele, suggesting that these alleles were in the repulsion phase (L982+F1534C and L982W+F1534) (Fig. 1*C*). Assuming that there is no concomitant L982W+F1534C haplotype in the Hoa Kien population, most of the dead individuals are heterozygous (L982W+F1534/L982+F1534C) and most of the surviving individuals are homozygous of L982W+F1534. There was a strong correlation between homozygotic L982W genotype and phenotype (dead/survived) (*p* < 2.2 × 10^−16^ in Fisher’s exact test) where 97.6% (40/41) of the survived individuals were L982W+F1534 homozygotes, whereas only 2.7% (1/37) of the dead individuals had the same genotype (Fig. 1*C*). In the Hanoi 2 population, the frequency of L982W was 97.5%, resulting in the fact that most individuals were homozygous for this allele. Unlike the Hoa Kien population, there were L982W homozygotes with heterozygous or homozygous F1534C (i.e., WW/FC), indicating that the Hanoi 2 population contained concomitant L982W+F1534C haplotypes. The frequencies of mosquitoes having the WW/FC genotype in the surviving and dead groups were 40.9% (27/66) and 7.1% (1/14), respectively (Fig. 1*D*). The survival rate was much higher in the mosquito group that had at least one L982W+F1534C haplotype than the mosquito group that did not have concomitant L982W and F1534C mutations in the Hanoi 2 population (Fig. 1*D*). Statistical analysis was conducted using Bayesian logistic analysis to evaluate the contribution of L982W to the resistance. The contribution value of L982W was 0.903 (95% confidence interval (CI), 0.644–0.997), which was much higher than the values for V1016G (0.122 with 95% CI, 0.003–0.450) and F1534C (0.146 with 95% CI, 0.003–0.519) suggesting strong correlation of L982W mutation with permethrin resistance.

### Establishment of various *kdr* strains and determination of the whole *Vssc* coding sequences (CDSs)

Since genotyping studies of *Vssc* showed that Hoa Kien and Hanoi 2 populations contained various *Vssc* haplotypes, we next attempted to establish several isogenic strains with homozygous genotypes on all of the V*ssc* polymorphic loci (Fig. 2*A*, *B*, and *SI Appendix*, Fig. S1). Since the frequency of the V1016G allele was very low in the two Vietnamese populations, we also tried isolating three additional strains from a population collected from Singapore in 2016 (*SI Appendix*, Table S1). After the first selections, conducted on the basis of the four alleles, namely, L982W, S989P, V1016G, and F1534C, we sequenced full-length *Vssc* coding regions individually using a combination of the target capture method and the next-generation sequencing (NGS) (*SI Appendix*, Fig. S1). When any heterozygous polymorphism was found, we repeated the isolation process using the polymorphisms newly found. In HK-C, isolated from the Hoa Kien population that had the F1534C allele, two different haplotypes were found to be included. Two HK-C derived strains, FC66 and FC213 had S66F or L213F alleles, respectively (Fig. 2*B*). Similarly, in the HK-W isolated from the Hoa Kien population and having the L982W allele, two additional haplotypes were found. Two new strains isolated from KH-W were designated as FTW (L199F+A434T+L982W) and FWI (L199F+L982W+T1385I) (Fig. 2*B*). Three strains were established from Hanoi 2 population; GY (V1016G+D1763Y), PGC (S989P+V1016G+F1534C), and FTWC (L199F+A434T+L982W+F1534C). Three strains were isolated from the Singapore population: 1534C (F1534C), PG (S989P+V1016G), and PGG (S989P+V1016G+V1703G) (Fig. 2*B*). Eventually, a total of 1594 mosquitoes were individually genotyped in 12 isolation processes, resulting in the establishment of 10 resistant *kdr* strains (Fig. 2*B*).

**Figure 2.**
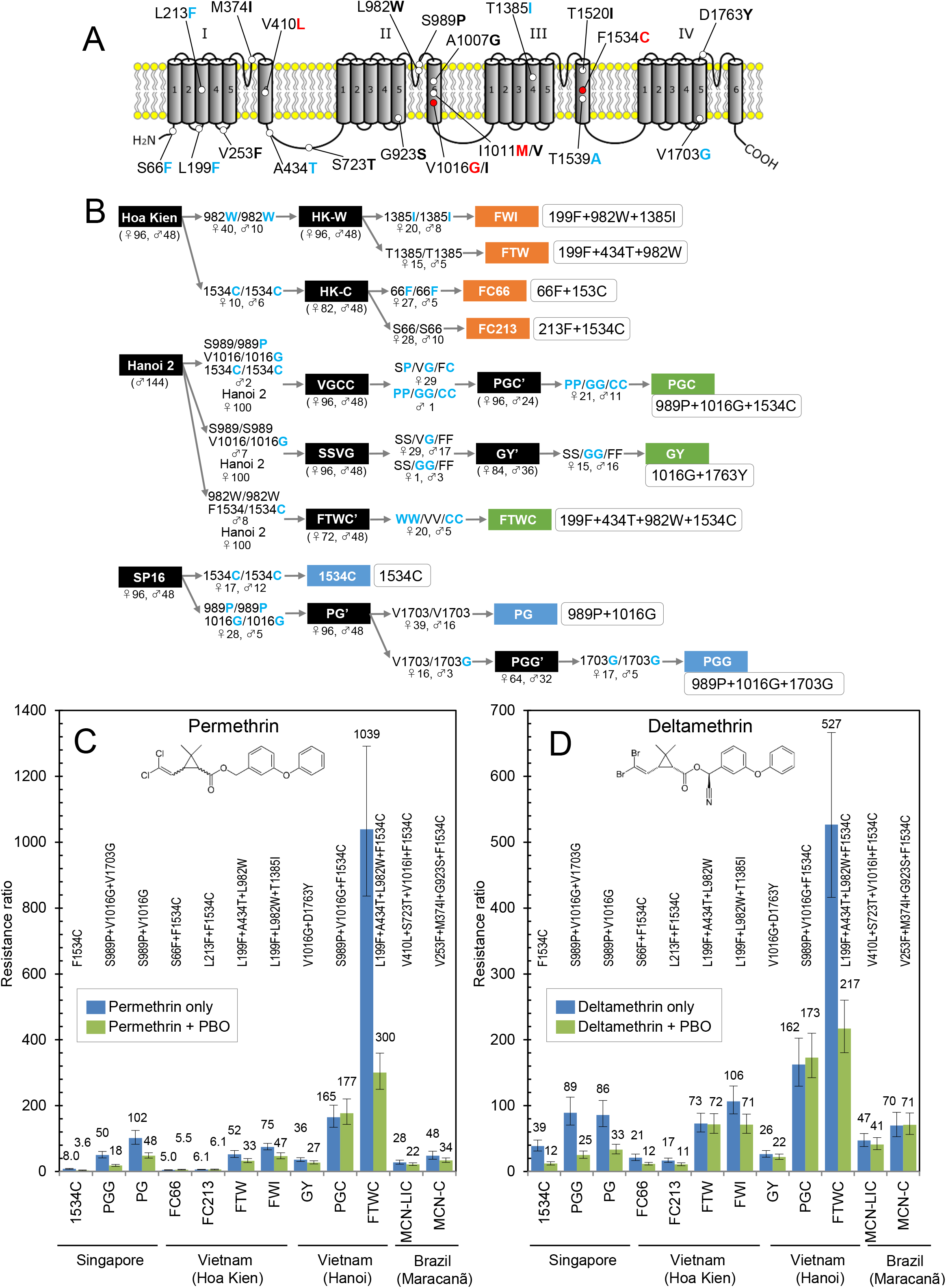
Establishment of 10 *kdr* strains of *Ae. aegypti* and their resistance levels to pyrethroids. (*A*) Amino acid substitutions found from *Vssc* of *Ae. aegypti*. Seven amino acid substitutions newly discovered in this study were indicated with blue letters. Three amino acid substitutions I1011M, V1016G, and F1534C have been proven their contribution to pyrethroid resistance by multiple electrophysiological studies (27). In this study, we numbered the amino acid position according to the sequence of the most abundant splice variant of the house fly *Vssc* (GenBank accession nos. AAB47604 and AAB47605). (*B*) Establishment of 10 *kdr* strains having different mutant haplotypes of *Vssc* homozygously. Substituted amino acids are indicated in blue bold. The numbers of mosquitoes genotyped were expressed in parentheses below population names. The homogeneity of isolated populations was verified via sequencing whole *Vssc* CDSs of at least eight mosquitoes individually via NGS (See Materials and Methods). (*C*, *D*) RRs of 12 *kdr* strains with different mutant *Vssc* haplotypes for two pyrethroid insecticides, permethrin (*C*) and deltamethrin (*D*). RRs were calculated by dividing the LD_50_ of each strain by the LD_50_ of the susceptible SMK strain. Error bars indicate the minimum and maximum 95% confidence interval (CI) for the RRs. RRs for SMK, MCNaeg-LIC, and MCNaeg-C are from our previous report (20).

### Susceptibilities of 10 *kdr* strains to pyrethroids

Ten resistant strains were tested for the susceptibility to two representative pyrethroids permethrin and deltamethrin (Fig. 2*C*, *D*, *SI Appendix*, Fig. S2). Two strains having L199F+L982W (FTW and FWI) showed approximately the same resistance level to permethrin (52 and 75 folds) and deltamethrin (73 and 106 folds), compared with the strains having V1016G allele; resistance ratios (RRs) of GY, PG, and PGG to permethrin and deltamethrin were 36–102 and 26–89 folds, respectively. FTWC (L199F+A434T+L982W+F1534C) strain showed the incomparably highest resistance level, with RRs of 1,039 and 527 folds, to permethrin and deltamethrin, respectively (Fig. 2*C* and *D*). Fifty percent lethal dose (LD_50_) of FTWC to permethrin was 130–208 times higher than that of three strains having F1534C (1534C, FC66, and FC213). The PGC (S989P+V1016G+F1534C) strain showed the second-highest resistance level, with RRs to permethrin and deltamethrin both being approximately 160-folds. Mosquitoes were treated with piperonyl butoxide (PBO) before applying pyrethroids to inhibit the detoxification of insecticide by cytochrome P450 monooxygenases. The synergistic ratios (SRs) of PBO on permethrin toxicity varied between strains; the SRs of FC66, FC213, FTW, and FWI, all isolated from the Hoa Kien population, were 2.4–4.2 (*SI Appendix*, Table S2). Conversely, among the three strains isolated from Hanoi 2 population, the SRs of PGC and GY were 2.5 and 3.5, respectively, and were not significantly different from those of the susceptible SMK (2.7). By contrast, the SR of FTWC to permethrin was 9.2, which is the highest value among the 11 strains tested (Fig. 2*C*, *SI Appendix*, Fig. S2, and Table S2). The RRs to permethrin under PBO-treated conditions were 3.6–6.1 folds in the three strains containing F1534C (1534C, FC66, and FC213) and 18–48 folds in the five strains having V1016G (GY, PG, and PGG). The RRs of 1531C and GY to permethrin+PBO were similar to those of *Ae. albopictus* having F1534C or V1016G alleles reported in previous studies (33). The RR of FTW (L199F+A434T+F982W) to permethrin with PBO was 33 folds, and these values increased to 300 folds with another amino acid substitution F1534C in FTWC strain. The RR of PG to permethrin was 48 folds, whereas that of PGC having F1534C allele was 177 folds under pretreatment of PBO. The RRs to deltamethrin+PBO were similar to those of permethrin, and the RRs of FTWC and PGC were 217 and 173 folds, respectively, which were 3.0 and 5.2 folds higher than those of FTW and PG, respectively (Fig. 2*D* and *SI Appendix*, Table S2). With PBO treatment, 1534C, FC66, and FC213 strains, having the F1534C allele, showed resistance to deltamethrin as well, and the RR of 11–12 folds was greater than that to permethrin (3.6–6.1 folds). The 95% CIs of RRs for deltamethrin in PGC and FTWC overlapped suggesting that resistance levels of these two strains to deltamethrin were not significantly different without detoxifications by P450 monooxygenases. The resistance levels of PGC and FTWC to permethrin and deltamethrin were much higher than those of MCN-LIC (V410L+S723T+V1016I+F1534C) and MCN-C (V253F+M374I+G923S+F1534C), which were established from a Brazilian population of *Ae. aegypti* (Fig. 2*C* and 2*D*) (20).

### DNA polymorphisms in *Vssc* CDSs of 10 *kdr* strains

The purities of the isolated strains were confirmed by sequencing the entire *Vssc* CDSs for at least eight individuals of each strain. Within 10 newly established strains and the susceptible SMK strain, 12 synonymous mutations and 27 nonsynonymous polymorphisms were found, compared with the reference LVP strain, which was used for the *Aedes* genome project (Fig. 3*A*) (34). Six of the 11 synonymous polymorphisms were newly found in this study (S66F, L199F, L213F, A434T, T1385I, and V1703G). Based on the polymorphisms each strain possesses, we predicted how each haplotype was created historically. It is estimated that *Vssc*s of both FTWC and PGC were generated not due to additive polymorphisms but due to crossing over events (Fig. 3*B*). A phylogenic tree created by the full-length CDSs of *Vssc* resulted in both haplotypes having V1016G and L982W fit into the single clades while F1534C distributed across multiple clades (Fig. 3*C*).

**Figure 3.**
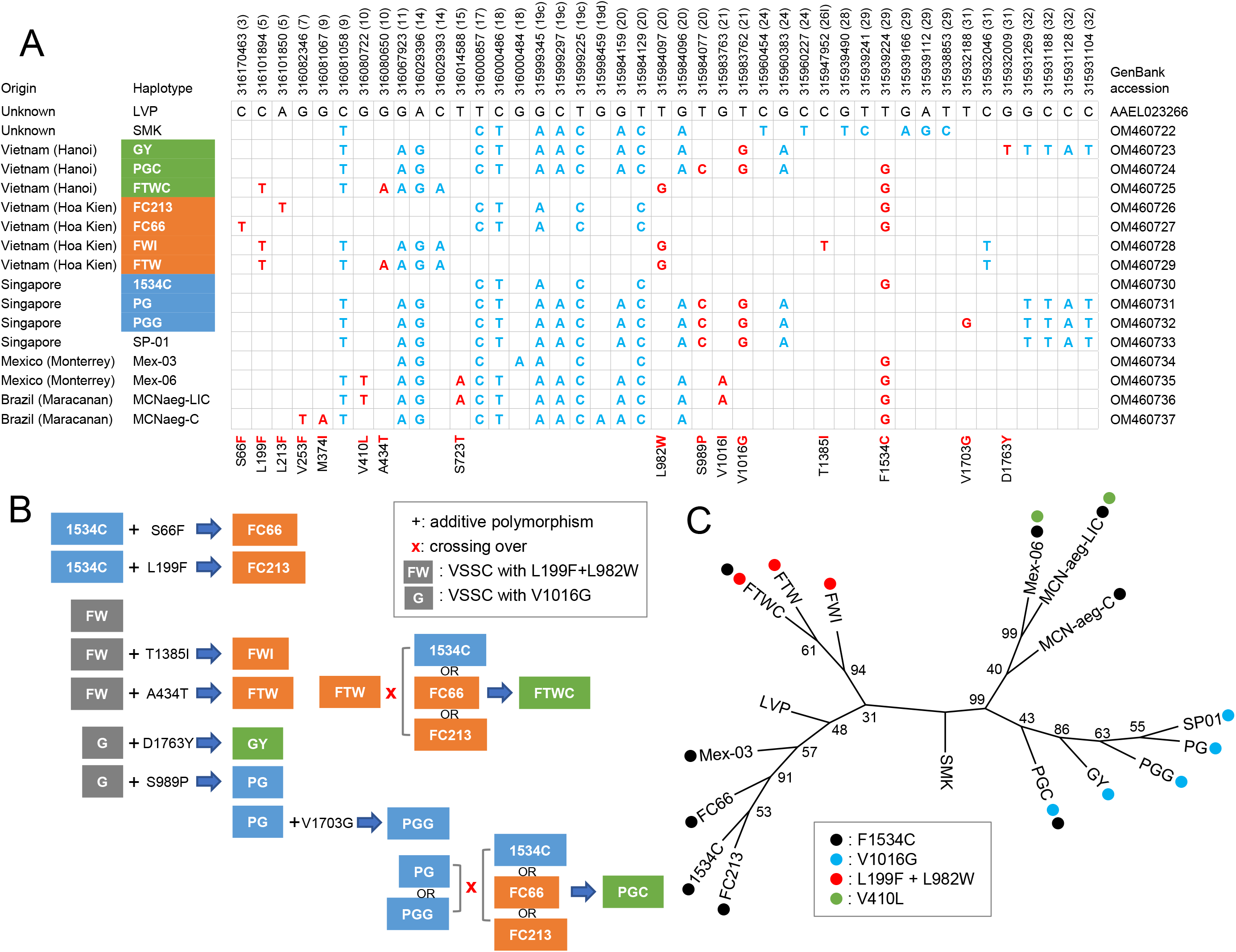
Polymorphisms and amino acid substitutions in the coding region of *Vssc* gene of *Ae. aegypti* strains. (*A*) All synonymous and nonsynonymous nucleotide polymorphisms found in entire CDSs of *Vssc* genes of 10 kdr strains. Data of two insecticide-susceptible strains, LVP and SMK, and five resistant mosquitoes from our previous report are also included (20). Accession for LVP is VectorBase accession number. (*B*) Expected evolutionary processes of *Vssc* genes obtained in this study. (*C*) Phylogenetic tree constructed by Genetyx software using the full-length CDSs of *Vssc* gene. Numbers on branches are the 1000 bootstrap values calculated using MEGA7.

### *Vssc* allele frequencies in *Ae. aegypti* field collected from Vietnam and Cambodia

To understand the allele frequency of *Vssc* in *Ae. aegypti* collected from Vietnam and Cambodia, 13 amino acids of *Vssc* were targeted and genotyped (Table 1). Another new amino acid substitution T1539A, which was found in the process of genotyping F1534C was also included (Fig. 2*A*). In this study, 368 field-collected nonbreeding insects (i.e., generation zero) were examined to understand the gene frequency more accurately. Five possible amino acid substitutions, L213F, V410L, V1007G, I1011M, and T1520I, were not detected in any of the field populations of *Ae. aegypti* collected from Vietnam (Hanoi, Dak Lak, and Ho Chi Minh) and Cambodia (Phnom Penh). The frequency of the L982W allele was high in Vietnam, accounting for more than 79% (Table 1, Fig. 4). Furthermore, the frequency of the concomitant mutations L982W+F1534C was estimated to be 28.4%–40.5%, 11.8%–16.5%, and 6.2%– 7.3% in the populations from Hanoi, Dak Lak, and Ho Chi Minh City, respectively. The frequency of V1016G, one of the major *kdr* alleles in Southeast Asia, was low in Vietnam, with 10.8% in Hanoi and <1% in Dak Lak and Ho Chi Minh (Fig. 4). Totally 51 individuals of *Ae. aegypti* collected from two water pools (larvae) and a vehicle (adults) in Phnom Penh, Cambodia, were genotyped. Since the genotype composition was quite simple and all genotypes were composed of any of five haplotypes, the frequencies of all haplotypes in the Phnom Penh populations could be estimated (*SI Appendix*, Fig. S3). The frequencies of L199F+A434T+L982W+F1534C and L199F+L982W+F1534C in the Phnom Penh population were 63.0% and 8.8%, respectively. Furthermore, the frequency of *Vssc* having S989P+V1016G+F1534C was 18.6%, resulting in >90% of the haplotypes in the Phnom Penh population were occupied by hyperresistant-type L982W+F1534C or V1016G+F1534C (Fig. 4 and *SI Appendix*, Fig. S3). The frequency of the L199F allele was also quite high in Vietnam (58.1%–98.9%) and Cambodia (78.4%). Of 368 mosquitoes tested, 266 were homozygous at the L199F allele and all were also having L982W allele homozygously, suggesting that L199F is strongly linked with L982W. Conversely, of 39 mosquitoes that had both L982W and F1534C homozygously, nine also had L199 homozygously, suggesting that L982W is not always accompanied by L199F when it is with F1534C (Table 1). The frequencies of A434T and T1385I in Vietnam ranged from 33.3% to 50.6% and from 4.2 to 28.1%, respectively. In this study, no amino acid substitution was detected at the I1011 allele, which was inconsistent with a previous study detecting I1011V all homozygously (30/30) in the populations collected at Nha Trang, Vietnam (35).

**Figure 4.**
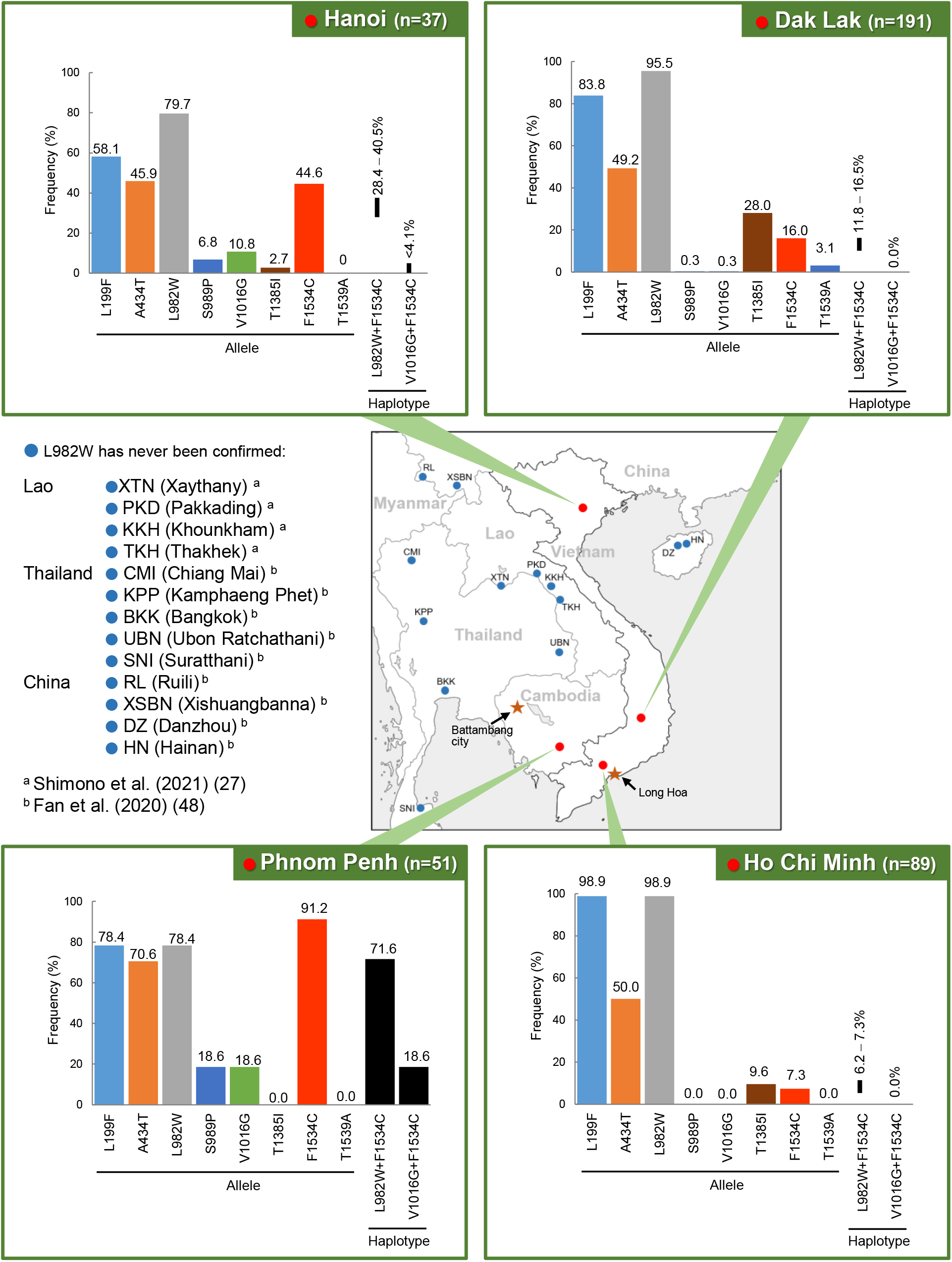
Hyper insecticide-resistant dengue vectors localizing in Indochina Peninsula. Frequencies of eight alleles and two haplotypes (L982W+F1534C and V1016G+F1534C) of *Vssc* in the field-collected *Ae. aegypti* collected from Vietnam and Cambodia are shown. Since the number of haplotypes is large, the exact frequency of two haplotypes cannot be calculated and minimum and maximum possible frequencies of these haplotypes are expressed for the populations from Vietnam. In the population from Phnom Penh, the exact frequency of each haplotype was estimated because of the restricted number of haplotypes. Please see Fig. S3 for the actual frequencies of all five haplotypes of *Vssc* in the populations from Phnom Penh. No L982W allele was detected from the cities marked with blue plots in China, Lao, Thailand, and Malaysia although the L982 allele was examined (26, 55). Note that L982W has never been detected in Indochina Peninsula except Vietnam and Cambodia.

**Table 1.**
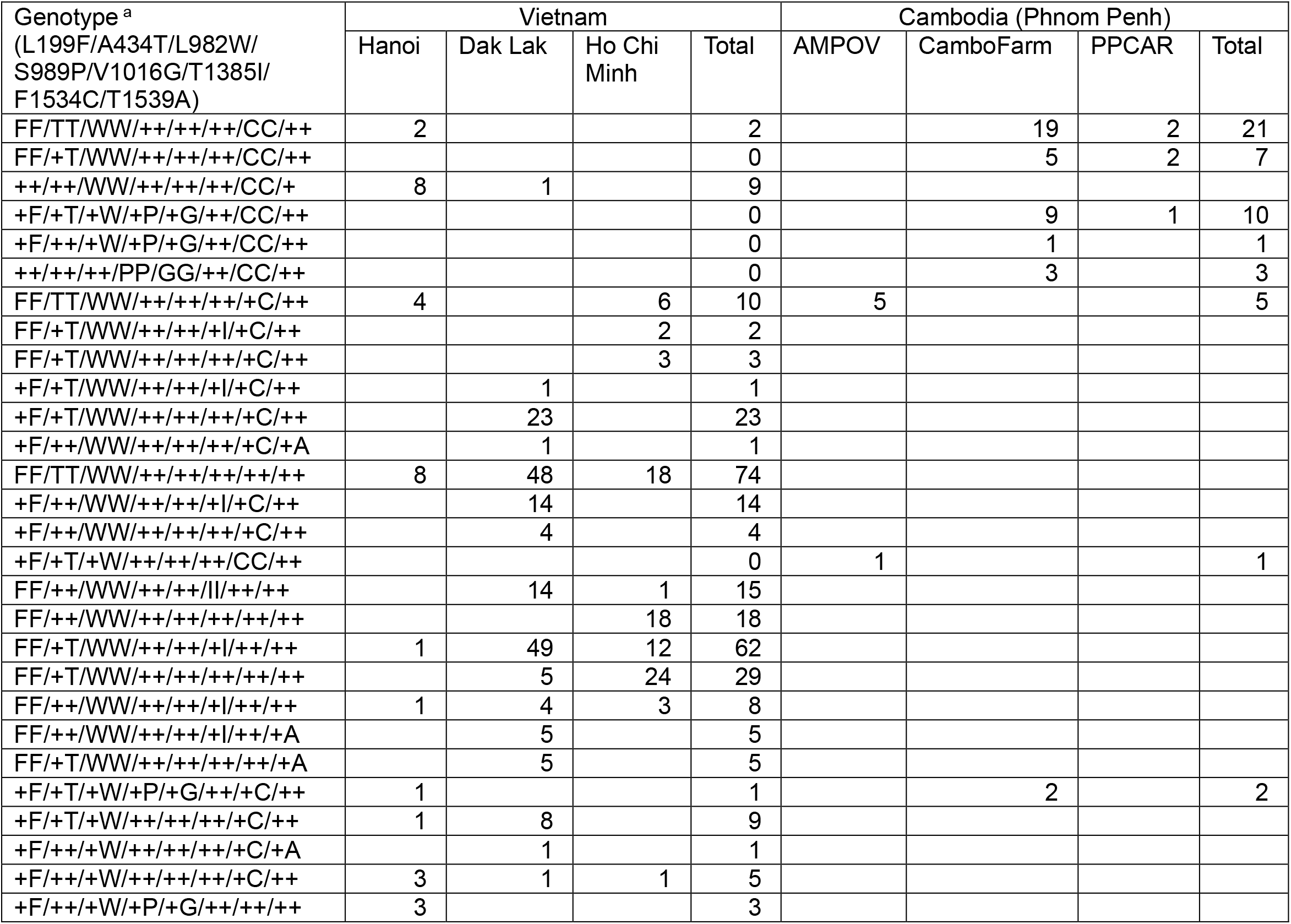
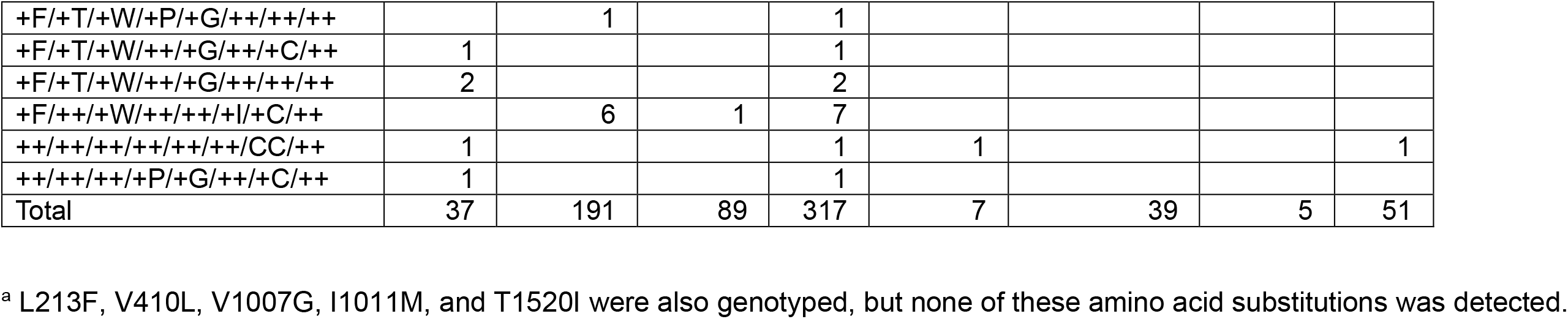
Genotype of *Vssc* in the field-collected (G0) *Ae. aegypti* from Vietnam and Cambodia

## Discussion

In the present study, the L982W allele located at the linker helix connecting S5 and S6 in domain II of *Vssc* was identified at a fairly high frequency in *Ae. aegypti* collected from Vietnam and Cambodia. L982W had been originally identified in an *Ae. aegypti* population collected before 1998 at Long Hoa, Vietnam (36). The contribution of this allele on pyrethroid resistance, however, has never been confirmed, and therefore, this allele has never been highlighted as a resistance factor. Chen et al. previously reported that the alanine to valine substitution in linker helix of the S5 and S6 in domain III of *Drosophila Vssc* conferred pyrethroid resistance (37). They proved this by electrophysiological studies using mutant cockroach *Vssc*s expressed in *Xenopus* oocytes. Remarkably, this amino acid position is topologically at the same position as L982W in domain II implying the important role of L982W in pyrethroid resistance (37). In this study, we established 10 *Ae. aegypti* strains that have various knockdown resistance alleles in *Vssc* and evaluated their resistance levels to pyrethroid insecticides. Strains having the L982W allele (FTW and FWI) showed similar or even higher levels of resistance to pyrethroids than those having V1016G (GY, PG, and PGG), a typical knockdown resistance allele in Asia. Concomitant mutations L982W+F1534C were also confirmed, and the strain that has these multiple *Vssc* mutations exhibited >1000 folds of resistance, which is >10 times higher resistance levels than any other field populations ever reported.

Kawada et al. conducted a nationwide survey of the pyrethroid susceptibility of *Ae. aegypti* between 2006 and 2008 in Vietnam (38). Larvae of *Ae. aegypti* collected from central and southern Vietnam were particularly resistant to *d*-T_80_-allethrin. The frequency of the V1016G allele was not so high throughout Vietnam, and the F1534C allele was identified at a certain frequency although no clear correlation with pyrethroid susceptibility was observed (19). It is possible that resistance of *Ae. aegypti* to *d*-T_80_-allethrin confirmed between 2006 and 2008 was mostly due to the L982W allele, which was not investigated at that genotyping study. In the present study, frequencies of L982W were higher at central (Dak Lak, 95.5%) and southern (Ho Chi Minh, 98.9%) populations (Fig. 4). We rechecked the raw data from the previous genotyping studies by Kawada et al. and calculated the frequency of the L982W allele in *Ae. aegypti* collected between 2006 and 2008 in Vietnam (19). Of 448 mosquitoes tested, 237 were homozygous and 50 were heterozygote for L982W, and the overall frequency of this allele was 58.5%. It is noteworthy that the frequency of L982W was lower than that of *Ae. aegypti* collected from Vietnam in 2016 and we could not confirm any individual having L982W+F1534C homozygously implying that this haplotype has been selected by pyrethroid treatments and increased its frequency after the year 2008.

L982W locates in the linker helix, connecting the fifth and sixth segments in the second domain of *Vssc* (Fig. 2*A*). E985 and E988 located in this region are called the inner and outer rings, respectively, and are playing very important roles for sodium–ion selectivity and permeation rate (39). We confirmed that the 31 amino acid sequences containing these two rings and L982 (PRW…FRV**L**_982_CG**E**_985_WI**E**_988_SMWDCM) are completely conserved among at least 80 insect species (*SI Appendix*, Fig. S4). Regarding the core sequence (**L**_982_CG**E**_985_WI**E**_988_), all seven amino acids of 80 insects are completely identical with *Vssc* of mammals such as rats, mice, and humans (*SI Appendix*, Fig. S4). It is presumed that this amino acid composition is highly important and essential to maintaining the function of *Vssc* beyond the borders of vertebrates and invertebrates. Both leucine and tryptophan are hydrophobic, nonpolar amino acids, but tryptophan has an aromatic ring, which may alter the intermolecular forces with ligands (*SI Appendix*, Fig. S5). L1014F, the first reported and the most common *kdr* allele in insects, is also the substitution from leucine (40). Phenylalanine is also a hydrophobic, nonpolar amino acid and has an aromatic ring-like tryptophan (*SI Appendix*, Fig. S5). Although the substitution from leucine to tryptophan may have had any costs for *Vssc* to maintain its function as a channel, the benefits may have outweighed the disadvantages in Vietnam and Cambodia, where strong selection pressure by pyrethroid insecticides is expected (41, 42).

To understand the effects of amino acid substitutions on the affinity of pyrethroids to the sodium channel, a homology model of *Ae. aegypti Vssc* was generated on the basis of electron microscopy crystal structure of the American cockroach *Periplaneta americana* (Linnaeus, 1758) NavPaS. The *Vssc* model with and without amino acid substitution(s) was subjected to docking simulation with 1R-*trans*-permethrin. The segments 5 and 6 of each domain conform pore domain, which is directly involved in ion permeation. Several *Vssc* alleles such as V1016G, L1014F, and F1534C, all of which have been proven for their association with pyrethroid resistance by electrophysiological studies, are located in these segments (11, 40). In this model, 1R-*trans*-permethrin was located at this binding pocket, so-called pyrethroid receptor site PyR1, as previously reported (Fig. 5*A*) (17). The molecular model exhibited that L982 locates at the region near V1016 and F1534 where permethrin interacts (Fig. 5*A* and *B*). The indolyl group of the aromatic ring of L982W, however, is a steric obstacle for permethrin for approaching this position (Fig. 5*C*). Furthermore, in L982W+F1534C, the degree of inhibition looks much stronger (Fig. 5*D*). Effects of L982W with F1534C on the relationship between permethrin and *Vssc* was quite similar to those of V1016G with F1534C (Fig. 5*E*–*G*). These results strongly support the fact that the FTWC strain having *Vssc* with L982W+F1534C exhibited a quite high level of pyrethroid resistance in the bioassay. Conversely, we conducted a similar docking simulation at the deduced second pyrethroid receptor PyR2 (17); however, this region has some distance with the L982 allele, and no significant effect on the interaction with permethrin was observed (data not shown).

**Figure 5.**
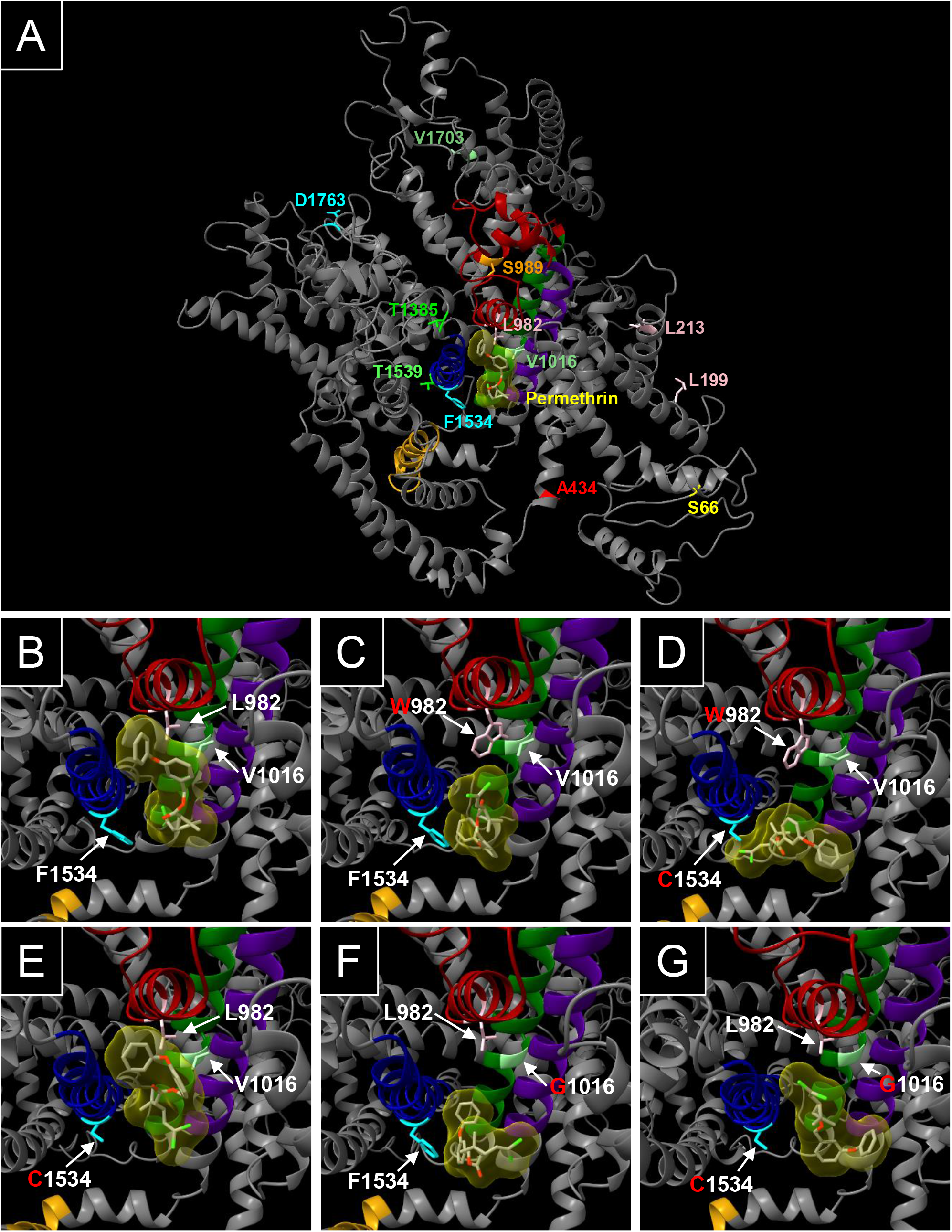
Docking simulation of *Ae. aegypti Vssc* (with and without *kdr* mutations) and 1R*-trans*-permethrin (yellow surface in the center of the channel). (*A*) Locations of the key amino acid residues L982, V1016, and F1534 and nine other amino acid residues found in this study. Note that the residues L982, V1016, and F1534 are surrounding permethrin configurating at the pyrethroid binding pocket. (*B*–*G*) Magnified models around pyrethroid binding pocket. (*B*) Wild type, nonmutated *Vssc*. (*C*) *Vssc* with L982W. (*D*) *Vssc* with concomitant L982W+F1534C. (*E*) *Vssc* with F1534C. (*F*) *Vssc* with V1016G. (*G*) *Vssc* with concomitant V1016G+F1534C. Note that concomitant substitutions L982W+F1534C severely interfere with permethrin approaching the normal binding position of *Vssc* (*D*) compared to the *Vssc* of wild type (*B*) and *Vssc* with a single amino acid substitutions L982W (*C*) or F1534C (*E*).

Novel amino acid substitutions have been identified by sequencing CDSs of *Vssc*: S66F, L199F, L213F, A434T, T1385I, T1539A, and V1703G. The combination of NGS and a concentration method of CDSs using target capture probes effectively discovered new SNPs as in previous studies (20, 43). Since all of these alleles locate at some distance from the permethrin binding region (Fig. 5*A*), and since there have been no reports of amino acid substitution at these locations from other resistant pest insects, these mutations are unlikely to affect the susceptibility of *Vssc* to pyrethroids. As for D1763Y, electrophysiological studies have ruled out its involvement in pyrethroid sensitivity (17). FTWC strain, which showed the highest level of resistance to pyrethroids among 10 resistant strains tested, had L199F and A434T alleles besides L982W and F1534C. L199F locates in the pore helix, which connects the second and the third segments of the first domain (Fig. 2*A*). This mutant allele is strongly linked with L982W since all mosquitoes having L199F homozygously had L982W homozygously (Table 1). Conversely, L982W is not always with L199F when it is accompanied by F1534C (Table 1). Although the role of L199F could not be derived clearly by molecular modeling analysis, it may sterically compensate for the disadvantage of *Vssc* as a sodium channel caused by the L982W mutation. To evaluate the effects of mutant alleles on pyrethroid resistance, the first generation (G1) of the three field populations of *Ae. aegypti*, Hanoi 2, Dak Lak 6, and HCM 4, were genotyped after selections with 59 ng permethrin. Thirteen amino acid residues that have been detected in this study or in previous reports were targeted (Fig. 2*A* and *SI Appendix*, Table S3). In the HCM 4 population, the estimated frequency of L199F+A434T+L982W (all homozygotes) in dead and surviving mosquitoes were 2.9% and 3.1%, respectively (*SI Appendix*, Table S3), whereas the estimated frequency of L199F+L982W (all homozygotes) in dead and surviving mosquitoes were 5.8% and 12.4%, respectively. These results seem to imply that A434T is not involved in the pyrethroid resistance. Unfortunately, all three populations examined included 25 genotypes resulting in the restricted number of each genotype, and we could not statistically evaluate the effect of each mutant allele (*SI Appendix*, Table S3). Conversely, 31/32 (96.9%) of the Hanoi 2 population having at least one L982W+F1534C haplotype survived, which is significantly higher than the survivorship (31/44, 70.5%) in the group having L982W homozygously but without F1534C, suggesting a strong effect of concomitant L982W+F1534C on permethrin resistance (*SI Appendix*, Table S3). Nevertheless, further studies are needed to elucidate the association of L199F and A434T on pyrethroid resistance using electrophysiological studies and/or gene editing technologies.

We have previously synthesized a *Vssc* full-length cDNA with the triple mutations S989P+V1016G+F1534C, expressed in the *Xenopus* oocytes, and measured its sensitivity to pyrethroids by electrophysiological studies (28). We observed that *Vssc* with these mutations was much less sensitive to both permethrin and deltamethrin than those with each single mutation. Soon after our report, *Ae. aegypti* having such triple-mutant *Vssc* alleles was reported from Myanmar, followed by various countries including Indonesia, Saudi Arabia, Sri Lanka, China, Thailand, Lao, and Malaysia (22-25, 44). In this study, we have established a strain (PGC) with triple amino acid substitutions and evaluated its susceptibility to pyrethroids *in vivo* for the first time. The PGC strain certainly exhibited a high level of resistance to both permethrin and deltamethrin than GY, PG, and PGG strains, all having V1016G but without F1534C, strongly supported our electrophysiological studies. Alternatively, multiple electrophysiological studies suggested that F1534C confers resistance to type I but not to type II pyrethroids (17, 28, 45), but in this study, 1534C strain as well as FC66 and FC213 strains showed more than 10-fold resistance to type II deltamethrin under the pretreatment of PBO. The remaining RRs may be due to any unknown resistance mechanism(s), but this (F1534C may confer resistance to type II pyrethroids) is consistent with the results of other reported studies (33, 46). Furthermore, our previous electrophysiological studies claimed that S989P alone does not confer permethrin susceptibility of *Vssc* but decreases deltamethrin sensitivity when it is combined with the V1016G allele (28). The RRs of GY (V1016G+D1703Y) and PG (S989P+V1016G) strains to deltamethrin (+PBO) were 22 folds (95% CI: 18.4–26.4) and 33 folds (95% CI: 26.9–41.3), respectively. Although the 95% CIs of these RRs do not overlap, no obvious differences were observed between the RRs of these two strains (Fig. 2*D*). The discrepancy between electrophysiological and *in vivo* studies was also mentioned in some recent studies (24, 47, 48), the reason for this has not been implicated though. Further study is needed to bridge the gaps between the results of electrophysiological studies and the actual levels of insecticide resistance.

There have been many reports of pyrethroid resistance in *Ae. aegypti* but relatively few in *Ae. albopictus* (11). In this study, we compared the permethrin susceptibilities between these two species collected from the water containers at the same areas and found that the level of resistance was clearly higher in *Ae. aegypti* (Fig. 1*B*). This is probably due to the differences in the ecology and habitats of the two species. Because of higher preference to humans as a blood-sucking source and high habitation affinity to humans, a higher percentage of *Ae. aegypti* are collected indoors than *Ae. albopictus* (49). In countries where dengue fever is endemic, vectors are mainly controlled by indoor pyrethroid fogging, so it is likely that adult *Ae. aegypti* have more chances of exposure to insecticides resulting in the development of resistance. Conversely, *kdr* genes, such as F1534C/S and V1016G, have also been recently detected from European and Asian populations of *Ae. albopictus*; thus, controlling this mosquito species by pyrethroids is also becoming difficult at some dengue epidemic regions (33, 50, 51).

PBO showed a strong synergistic effect on permethrin toxicity in the FTWC strain, suggesting that detoxification by cytochrome P450 monooxygenases is also an important resistance mechanism in addition to *kdr*. Moreover, it was found that mosquitoes can develop extremely high levels (>1,000 folds) of resistance when they possess multiple resistance factors. We have previously reported that SP strain originally collected from Singapore in 2009 and artificially selected by permethrin for 10 generations in the laboratory developed 1,650 folds of resistance (52). In this case, increased detoxification activity by cytochrome P450 oxidases was involved in the resistance besides the S989P+V1016G-type *kdr* (52). The RR of SP to permethrin under PBO treatment was 34 folds, which was similar to that of the PG (48 folds) in this study. Given that the synergistic effect of PBO on permethrin toxicity in the SP strain was 82, FTWC can potentially and theoretically develop >24,000 folds of resistance (300 × 82), if this strain possesses the same detoxification mechanisms as SP. Studies on *Anopheles gambiae* (Giles, 1902) and *Anopheles funestus* (Giles, 1900), the major malaria vectors, have also reported that increased cuticle thickness reduces the penetration of insecticides and causes resistance (53, 54). Since similar resistance mechanisms are possibly involved in the pyrethroid resistance of *Ae. aegypti*, it will be necessary to focus on such other mechanisms as well in the future.

F1534C does not confer a very high level of resistance by itself, but it causes a much stronger level of resistance when it occurs together with other amino acid substitutions such as L982W and V1016G. The analysis of the full-length CDSs of *Vssc* of two highly resistant strains FTWC and PGC established in this study can decipher that the multiple resistance alleles were accumulated by recombination events rather than by sequential mutations (Fig. 3*A* and *B*). In FTWC and PGC, 3′ region of *Vssc* CDSs have a polymorphism that causes F1534C, but both genes also lose some polymorphisms(s) that are characteristic to FTW and PG/PGG/GY, supporting that these genes were produced by crossing over events of multiple genes (Fig. 3*A* and *B*). It is strongly suggested that genetic recombination is an efficient evolutionary means for insects to develop a high level of insecticide resistance, and thus, it was the driving force behind the rapid development of resistance in *Ae. aegypti*. There is no denying the possibility that the hyper resistant haplotypes of *Vssc* found in this study will be further recombined with each other to produce even stronger resistance genes.

The L982W allele is located near S989P and V1016G on the *Vssc*, and all can be genotyped by sequencing the same PCR product of domain II. It is a sort of mystery that the L982W allele has never been identified in the recent studies of *Ae. aegypti* from 12 regions of the neighboring countries: Thailand, Lao, and China although this allele was focused (Fig. 4) (26, 55). L982W allele was also not detected from 442 *Ae. aegypti* collected from three regions of Nepal in 2017 and 2018 (56). Thus far, there is no information regarding the fitness cost of the L982W allele on the viability of mosquitoes, which we must investigate; at least in the laboratory, we do not experience difficulties in breeding FTWC strain compared with other *Ae. aegypti* strains. It is thus imaginable that populations carrying these mutant alleles will expand and spread to the whole Indochina Peninsula and from Southeast Asia to other tropical and subtropical regions of the world in the future. A case in point is the finding of this important haplotype L982W+F1534C in an *Ae. aegypti* population that invaded and bred at an international airport of Japan (57). Fortunately, the population was not sustained in the temperate climate. The expansion of these concomitant mutant alleles may face less resistance in tropical and subtropical countries and would hamper the control of *Ae. aegypti* and the diseases it vectors. Particularly in Phnom Penh city, two haplotypes (V1016G+F1534C and L199F+L982W+F1534C) that confer hyper resistance to pyrethroids are expected to account for most of the population, making it nearly impossible to control *Ae. aegypti* with pyrethroids in this region. This is also consistent with the fact that *Ae. aegypti* collected in Phnom Penh showed an extremely high level of resistance to many pyrethroid insecticides (32). The only previous report on the genotyping of *Vssc* in *Ae. aegypti* population from Cambodia showed that all 10 individuals collected at Battambang city, 280 km away from Phnom Penh, had F1534C with wild-type V1016 all homozygously (41). Country-wide survey of the knockdown resistance gene in *Ae. aegypti* is required in Cambodia.

## Conclusion

Our studies have shown that concomitant L982W+F1534C and V1016G+F1534C of *Vssc* confer an eminently higher level of pyrethroid resistance than the previously reported single *kdr* alleles F1534C and V1016G. Here, we emphasize strengthening the monitoring of these mutant alleles, especially in South East Asia to take countermeasures before they spread globally.

## Materials and Methods

### Mosquitoes

The larvae of *Ae. aegypti* were collected from fields of four countries listed in *SI Appendix*, Table S1. The larvae were reared in the laboratory as previously described (52). Species of mosquitoes were identified in the adult stage on the basis of the keys followed by bleeding in the laboratory.

### Bioassays

Simplified bioassays were conducted to evaluate the susceptibility of adult female *Ae. aegypti* to a representative pyrethroid insecticide, permethrin (91.2%, Sumitomo Chemical Co., Ltd., Osaka, Japan). Pools of 20 respective adult mosquitoes (4 to 6-days old) were treated in four replicates with each permethrin dose (5.9 and 59 ng) by topical applications and mortalities were assessed as described previously (33); 5.9 ng permethrin is equivalent to LD_99_ of a pyrethroid susceptible strain of *Ae. albopictus* (33). After the establishment of 10 *kdr* strains based on *Vssc* mutant alleles, additional bioassays were conducted using permethrin and deltamethrin (99.4%, GL Sciences Inc., Tokyo, Japan) to compare the pyrethroid susceptibility among strains as previously described (33). LD_50_s were calculated using Finney’s log-probit mortality regression analysis (58), implemented in R version 3.3.3 (www.r-project.org). RRs were calculated as the LD_50_ of each strain by the LD_50_ of the susceptible SMK strain. The 95% CIs of RR were determined by calculating the RRs for the minimum and maximum 95% CIs of LD_50_ values (46). Two RR values were considered significantly different if the minimum and maximum RR values did not overlap (46). The synergism of PBO (98.0%, Wako Chemical Pure Industries, Ltd., Osaka, Japan) was examined to evaluate the contribution of detoxification by cytochrome P450 monooxygenases (52).

### Genotyping

Genomic DNA was extracted from the legs of mosquitoes as previously described (33) and the partial genomic DNA of *Vssc* was sequenced individually. To examine the effect of known *kdr* alleles, S989P, V1016G, and F1534C were genotyped for dead and survived mosquitoes from two highly resistant Hoa Kien (n = 78) and Hanoi 2 (n = 80) populations used for the bioassay. After sequencing the whole CDSs of *Vssc* from established strains as described below, 13 *Vssc* alleles were targeted for the genotyping studies; L199F, L213F, V410L, A434T, L982W, S989P, A1007G, I1011M/V, V1016G/I, T1385I, T1520I, F1534C, and T1539A (Fig. 2*A* and *SI Appendix*, Fig. S6–S12). The A1007G allele was reported in the previous report (59). The condition of the PCR and the sequencing procedure were described previously (52). The PCR elongation time was differentiated according to the length of the PCR products (60 s/kbp). Primers used for PCR and sequencing were listed in the *SI Appendix*, Table S4.

### Mosquito selections by *Vssc* genotypes and establishment of *kdr* strains

Since genotyping studies of *Vssc* showed that Hoa Kien and Hanoi 2 populations contained various *Vssc* haplotypes, we attempted to establish several isogenic strains with homozygous genotypes on all of the V*ssc* polymorphic loci. Since the frequency of the V1016G allele was very low in the two Vietnamese populations, we also tried isolating three additional strains from a population collected from Singapore in 2016 (Fig. 2*B* and *SI Appendix*, Fig. S1) (33). In this study, we numbered the amino acid position according to the sequence of the most abundant splice variant of the house fly *Vssc* (GenBank accession numbers AAB47604 and AAB47605). Four (FC66, FC213, FTW, and FWI) and three strains (1534C, PG, and PGG) were established from Hoa Kien and Hanoi 2 populations, respectively. Three strains (1534C, PG, and PGG) were established from the Singapore SP strain. According to preliminary genotyping studies, a very low frequency of L982W+F1534C was found in the Hanoi 2 population; thus, 144 males were individually genotyped for L982W, S989P, V1016G, and F1534C using genomic DNA from a single hind leg. The small numbers of males having the same *Vssc* genotypes (L982W/F1534C: WW/FC, S989P/V1016G/F1534C: SP/VG/CC, and S989P/V1016G: SS/VG) were each mated with 100 virgin females of Hanoi 2 population and offspring were utilized for additional isolating steps (Fig. 2*B*). Eventually, three new *kdr* strains (PGC, GY, and FTWC) were established (Fig. 2*B*). Bioassays for permethrin and deltamethrin were conducted as described above and previously (52) for 10 resistant strains to obtain LD_50_s.

### Sequencing the whole coding region of the *Vssc* gene

Genomic DNA was extracted from individual mosquitoes using the MagExtractor Genome kit (Toyobo Co., Ltd., Osaka, Japan) (60). Index library construction and targeted capture of pooled libraries were conducted as previously described (60). Illumina library construction and hybridization capture was conducted with the biotinylated oligo probe designed from the *Ae. albopictus Vssc* gene, whose exons show >92.5% homologies to the exons in *Ae. aegypti* (60). The quantified library was sequenced using the Illumina MiniSeq with the Mid Output Kit (Illumina, Inc., San Diego, CA) and 151 cycles for both ends. Read pairs of 114– 349 kbp were sequenced for each sample. Raw fastq read data were deposited to the National Center for Biotechnology Information (NCBI) Sequence Read Archive (BioProject ID: PRJNA795523). NGS-based reads were mapped to the reference *Vssc* genome sequence (AALF000723-RA) and annotated for the synonymous and nonsynonymous nucleotide polymorphisms by using the automated MoNas pipeline (https://github.com/ItokawaK/MoNaS) as previously described (60).

### Evolutionary tree

The evolutionary history was inferred using the UPGMA method (61). The percentage of replicate trees in which the associated taxa clustered together in the bootstrap test (1,000 replicates) are shown (62). The evolutionary distances were computed using the Kimura two-parameter method (63) and are in the units of the number of base substitutions per site. The rate variation among sites was modeled with a gamma distribution (shape parameter = 1). This analysis involved 17 nucleotide sequences (*SI Appendix*, Table S5). All ambiguous positions were removed for each sequence pair (pairwise deletion option). There were 6709 positions in the final dataset. Evolutionary analyses were conducted in MEGA X (64). A phylogenic tree was constructed by GENETYX software (Ver 13, GENETYX Corp., Tokyo, Japan).

### Generation of homology model and ligand docking

Homology model of *Ae. aegypti Vssc* (Vectorbase accession: AAEL023266) was generated on the basis of the electron microscopy crystal structure of the voltage-sensitive sodium channel NavPaS from the American cockroach *Periplaneta americana* (PDB accession number: 6A90, DOI:10.2210/pdb6A90/pdb) (65). The model was produced using the Modeller Ver10.1 program (66). Automated docking of 1R-*trans*-permethrin (PDB accession number: 40159) with the model of the sodium channel was performed using the AutoDock 4.2.6 software package (67). Docking simulations were performed for 100 runs for each trial to generate histograms of the binding energies of each run. The root-mean-square deviation tolerance was set at 2 Å in the simulations.

### Statistical analysis

The association of *kdr* mutations and pyrethroid resistance was analyzed via Bayesian logistic analysis using Cmdstan Ver. 2.28.2 in R Ver. 4.1.2 (https://mc-stan.org). The model formula was yi~Be(logit(Pi)) and Pi = a + b1×1i + b2×2i + b3×3i + b4×4i., where Be represented Bernoulli distribution; i represented Individual Number; yi represented alive(1) or dead(0), x1 to x4 represented Input(x1 = Hoa Kien(0) or Hanoi(1), x2 = L982(0) or W982(1), x2 = V1016(0) or G1016(1), x3 = L1534(0) or C1534(1)); a represented intercept, and b1, b2, b3, and b4 represented parameters (the ranges were defined 0–1).

## Data availability

The short reads obtained via the NGS analysis for the 12 strains of *Ae. aegypti* are available in the NCBI Sequence Read Archive in BioProject PRJNA795523 at https://www.ncbi.nlm.nih.gov/bioproject/795523. Accession numbers of full-length amino acid sequences for 11 strains of *Ae. aegypti* are shown in *SI Appendix*, Table S5. All other study data are included in the article and/or *SI Appendix*.

## Acknowledgments

We thank Takashi Tsunoda for his support in mosquito collection. We are also grateful to Chigusa Yosida for her technical assistance in mosquito maintenance. This research was partially supported by AMED under grant Numbers JP17fm0108018, JP20fk0108067, JP20wm0225007, JP21wm0125006, JP21wm0225007, and JP21fk0108613.

