## Supplementary information for "Emergence of hyper insecticide-resistant dengue vectors in Indochina Peninsula: threats of concomitant knockdown resistance mutations"

**This PDF file includes:**
Table S1 to S5
Fig. S1 to S12

Table S1

Information of *Aedes aegypti* populations used for bioassays and genotyping studies

| Population name in Fig. 1 | Collection site | Code name | Number of water pools | Latitude | Longitude | Collection date | Generation tested |
|---|---|---|---|---|---|---|---|
| SMK[a] | Unknown | — | — | — | — | — | — |
| Hanoi 2 | Tu Liem, Hanoi, Vietnam | HNI-PK6-9 | 4 | 21.0 | 105.8 | July 17, 2016 | G1 |
| Hanoi 5 | Tu Liem, Hanoi, Vietnam | HNI-TL2, 4 | 2 | 21.0 | 105.8 | Nov 11, 2016 | G2 |
| Hanoi 6 | Tu Liem, Hanoi, Vietnam | HNI-H3, 5 | 2 | 21.0 | 105.8 | Feb 02, 2016 | G2 |
| Hoa Kien | Hoa Kien, Phu Yen, Vietnam | Hoa Kien | 21 | 13.1 | 109.2 | Feb 20, 2016 | G1 |
| Dak Lak 1 | Yok Don National Park, Dak Lak, Vietnam | DL-P1, 2 | 2 | 12.7 | 107.7 | Sep 22, 2016 | G2 |
| Dak Lak 3 | Buon Ma Thuot, Dak Lak, Vietnam | DL-U1-1, 2, 3, 4, 5 | 5 | 12.7 | 108.0 | Sep 19, 2016 | G1 |
| Dak Lak 5 | Buon Ma Thuot, Dak Lak, Vietnam | DL-U1-6, 7, 8, 10 | 4 | 12.7 | 108.0 | Sep 19, 2016 | G1 |
| Dak Lak 6 | Buon Ma Thuot, Dak Lak, Vietnam | DL-U1-11 | 1 | 12.7 | 108.0 | Sep 19, 2016 | G1 |
| Dak Lak 7 | Buon Ma Thuot, Dak Lak, Vietnam | DL-U2-4, 5, ex2, ex3 | 4 | 12.7 | 108.0 | Sep 20, 2016 | G1 |
| Dak Lak 8 | Yok Don National Park, Dak Lak, Vientam | DL-F2-1, F2-4-1, F2-8 | 3 | 12.7 | 107.7 | Sep 12, 2016 | G1 |

| | | | | | | | |
|---|---|---|---|---|---|---|---|
| Dak Lak 9 | Ho Chi Minh City, Vietnam | HCM108-2 | 1 | 10.8 | 106.7 | Sep 20, 2016 | G2 |
| Ho Chi Minh 4 | Ho Chi Minh City, Vietnam | HCM123-1, 2, 3, 4 | 4 | 10.8 | 106.7 | Sep 13, 2016 | G1 |
| Ho Chi Minh 6 | Ho Chi MinhCity, Vietnam | HCM-123-6 | 1 | 10.8 | 106.7 | Sep 13, 2016 | G1 |
| Mangrove Park | Surabaya City, East Java, Indonesia | Mangrove Park | 5 | -7.31 | 112.8 | Apr 19, 2018 | G2 |
| Lomboku | Pulau Lombok, Indonesia | Lomboku | 1 | -8.56 | 116.1 | Apr 23, 2018 | G2 |
| Petemon Kali 2 | Surabaya City, East Java, Indonesia | Petemon Kali 2 | 10 | -7.26 | 112.7 | Apr 18, 2018 | G2 |
| Simo Sidomulyo | | Simo Sidomulyo | 10 | -7.25 | 112.7 | Apr 17, 2018 | G2 |
| Pingtung | Pingtung City, Taiwan | Pingtung | 2 | 22.6 | 120.6 | Mar 08, 2016 | G2 |
| Kaohsiung | Kaohsiung City, Taiwan | Kaohsiung | 1 | 22.6 | 120.3 | Mar 2016 | G2 |
| Tainan | Tainan City, Taiwan | Tainan | 1 | 23.0 | 120.2 | Mar 03, 2016 | G2 |
| Aburi | Aburi, Ghana | Aburi | 8 | 5.85 | -0.123 | Jan 29, 2018 | G2 |
| Labadi | Labadi, Accra, Ghana | Labadi | 5 | 5.57 | -0.157 | Feb 02, 2018 | G2 |
| Maracanã | Maracanã, Rio de Janeiro, Brazil | Maracanã | 1 | -22.9 | -43.2 | Mar 13, 2016 | G2 |
| AMPOV | Ampov Prey Village, Phnom Penh, Cambodia | Ampov | 1 | 11.4 | 104.9 | May 29, 2019 | G0 |

| | | | | | | | |
|---|---|---|---|---|---|---|---|
| CamboFarm | Royal University of Agriculture, Phnom Penh, Cambodia | RUA_farm | 1 | 11.5 | 104.9 | May 30, 2019 | G0 |
| PPCAR | Collected in a vehicle by net sweeping at Phnom Penh, Cambodia | PP_car | 1 (adults collected in a car) | | | May 29&30, 2019 | G0 |
| Singapore | Singapore | SP16aeg | | 1.4 | 103.9 | 2016 | |

[a] SMK: Insecticide susceptible strain (1).

Table S2

Toxicity of permethrin and deltamethrin against eleven strains of *Aedes aegypti* having different voltage-sensitive sodium channel alleles

| Insecticide | Strain | n[a] | Slope ± SE[b] | $LD_{50}$ (95% CI[c]) | Synergistic ratio (SR[d]) | Ratio of SR (SRR[e]) | Resistance ratio (95% CI[f]) |
|---|---|---|---|---|---|---|---|
| Permethrin only | SMK[g] | 246 | 6.0±0.57 | 1.54 (1.44–1.65) | — | — | — |
| | 1534C | 460 | 3.5±0.28 | 12.2 (11.2–13.4) | — | — | 7.9 (6.8–9.3) |
| | FC213 | 320 | 4.8±0.53 | 9.4 (8.7–10.1) | — | — | 6.1 (5.3–7.0) |
| | FC66 | 380 | 4.2±0.40 | 7.7 (7.1–8.3) | — | — | 5.0 (4.3–5.8) |
| | GY | 200 | 5.1±0.48 | 55.4 (50.9–60.4) | — | — | 36 (31–42) |
| | PGG | 240 | 3.3±0.41 | 77.6 (68.6–87.8) | — | — | 50 (42–61) |
| | PG | 260 | 3.5±0.35 | 157 (137–180) | — | — | 102 (83–125) |
| | PGC | 567 | 2.1±0.16 | 254 (222–290) | — | — | 165 (135–201) |
| | FWI | 290 | 6.4±0.65 | 115 (108–123) | — | — | 75 (66–85) |
| | FTW | 200 | 3.7±0.45 | 80.3 (70.4–91.5) | — | — | 52 (43–64) |
| | FTWC | 487 | 2.1±0.21 | 1600 (1380–1860) | — | — | 1040 (836–1290) |
| | MCNaeg-LIC[h] | 340 | 2.5±0.25 | 42.9 (36.7–50.2) | — | — | 28 (22–35) |
| | MCNaeg-C[i] | 340 | 1.9±0.19 | 73.7 (61.4–88.5) | — | — | 48 (37–61) |
| Permethrin +PBO | SMK[g] | 322 | 5.1±0.47 | 0.576 (0.531–0.625) | 2.7 | 1.0 | — |
| | 1534C | 200 | 3.5±0.28 | 2.09 (1.84–2.37) | 5.8 | 2.2 | 3.6 (2.9–4.5) |
| | FC213 | 264 | 3.7±0.41 | 3.52 (3.17–3.92) | 2.7 | 1.0 | 6.1 (5.1–7.4) |
| | FC66 | 242 | 4.6±0.51 | 3.17 (2.86–3.52) | 2.4 | 0.9 | 5.5 (4.6–6.6) |

| | | | | | | | |
|---|---|---|---|---|---|---|---|
| | GY | 370 | 4.2±0.37 | 15.8 (14.5–17.3) | 3.5 | 1.3 | 27 (23–33) |
| | PGG | 647 | 3.7±0.26 | 10.6 (9.81–11.5) | 7.5 | 2.8 | 18 (16–22) |
| | PG | 320 | 4.3±0.42 | 27.7 (25.3–30.3) | 5.7 | 2.1 | 48 (40–57) |
| | PGC | 300 | 2.9±0.32 | 102 (89.6–117) | 2.5 | 0.9 | 177 (143–220) |
| | FWI | 278 | 3.9±0.47 | 27.1 (24.2–30.3) | 4.2 | 1.6 | 47 (39–57) |
| | FTW | 240 | 3.8±0.48 | 18.9 (17.0–21.0) | 4.2 | 1.6 | 33 (27–40) |
| | FTWC | 245 | 3.7±0.46 | 173 (156–191) | 9.2 | 3.5 | 300 (250–360) |
| | MCNaeg-LIC[h] | 400 | 4.3±0.44 | 12.8 (11.5–14.1) | 3.4 | 1.3 | 22 (18–27) |
| | MCNaeg-C[i] | 400 | 3.5±0.34 | 19.5 (17.4–21.8) | 3.8 | 1.4 | 34 (28–41) |
| Deltamethrin | SMK[g] | 529 | 5.0±0.42 | 0.0788 (0.0711–0.0872) | — | — | — |
| | 1534C | 240 | 3.7±0.46 | 3.05 (2.73–3.41) | — | — | 39 (31–48) |
| | FC213 | 400 | 5.0±0.42 | 1.32 (1.22–1.45) | — | — | 17 (14–20) |
| | FC66 | 240 | 3.5±0.39 | 1.66 (1.45–1.89) | — | — | 21 (16–27) |
| | GY | 280 | 5.5±0.53 | 2.08 (1.90–2.27) | — | — | 26 (22–32) |
| | PGG | 343 | 2.7±0.27 | 7.03 (6.16–8.02) | — | — | 89 (71–113) |
| | PG | 280 | 3.2±0.33 | 6.76 (5.96–7.66) | — | — | 86 (68–108) |
| | PGC | 615 | 2.7±0.24 | 12.8 (11.3–14.4) | — | — | 162 (130–203) |
| | FWI | 509 | 3.0±0.24 | 8.39 (7.63–9.23) | — | — | 107 (88–130) |
| | FTW | 485 | 3.7±0.31 | 5.74 (5.23–6.30) | — | — | 73 (60–89) |
| | FTWC | 360 | 2.7±0.24 | 41.5 (36.3–47.4) | — | — | 527 (416–667) |
| | MCNaeg-LIC[h] | 320 | 4.4±0.51 | 3.6 (3.3–4.1) | — | — | 47 (38–58) |
| | MCNaeg-C[i] | 400 | 2.2±0.24 | 5.4 (4.6–6.4) | — | — | 70 (53–90) |

| | | | | | | | |
|---|---|---|---|---|---|---|---|
| Deltamethrin +PBO | SMK[g] | 330 | 3.8±0.33 | 0.0230 (0.0208–0.0255) | 3.4 | 1.0 | — |
| | 1534C | 398 | 3.6±0.34 | 0.279 (0.253–0.308) | 10.9 | 3.2 | 12 (9.9–15) |
| | FC213 | 207 | 5.2±0.60 | 0.248 (0.225–0.274) | 5.3 | 1.6 | 11 (8.8–13) |
| | FC66 | 326 | 5.1±0.51 | 0.268 (0.248–0.288) | 6.2 | 1.8 | 12 (9.7–14) |
| | GY | 562 | 3.3±0.29 | 0.507 (0.469–0.549) | 4.1 | 1.2 | 22 (18–26) |
| | PGG | 280 | 3.4±0.35 | 0.574 (0.509–0.647) | 12.2 | 3.6 | 25 (20–31) |
| | PG | 240 | 3.9±0.43 | 0.768 (0.686–0.859) | 8.8 | 2.6 | 33 (27–41) |
| | PGC | 382 | 3.8±0.32 | 3.98 (3.63–4.37) | 3.2 | 0.9 | 173 (142–210) |
| | FWI | 287 | 4.6±0.43 | 1.64 (1.48–1.81) | 5.1 | 1.0 | 71 (58–87) |
| | FTW | 200 | 4.7±0.59 | 1.65 (1.48–1.83) | 3.5 | 1.5 | 72 (58–88) |
| | FTWC | 280 | 5.0±0.50 | 4.99 (4.60–5.41) | 8.3 | 2.4 | 217 (180–260) |
| | MCNaeg-LIC[h] | 520 | 3.9±0.43 | 0.95 (0.85–1.07) | 3.8 | 1.1 | 41 (33–51) |
| | MCNaeg-C[i] | 400 | 3.7±0.42 | 1.63 (1.44–1.85) | 3.3 | 1.0 | 71 (56–89) |

[a] n: number of females used for bioassay.

[b] SE: standard error.

[c] CI: confidence interval.

[d] SR: synergistic ratio = $LD_{50}$ (without PBO) / LD50 (with PBO).

[e] SRR: SR ratio = (SR of each strain) / (SR of SMK).

[f] CI of Resistance ratios were determined by calculating the resistance ratios for the minimum and maximum $LD_{50}$ values based on the $LD_{50}$ 95% CI.

[g, h, i] Data from our previous studies (2).

Table S3

Associations between permethrin susceptibility and *Vssc* genotypes in *Aedes aegypti* collected from Vietnam

| Genotype (L199F/A434T/L982W/ S989P/V1016G/T1385I/ F1534C/T1539A)[a] | Hanoi 2 G1[b] (mortality[c]: 17.5%) | | | | Dak Lak 6 G1[b] (mortality[c]: 30.0%) | | | | HCM 4 G1[b] (mortality[c]: 28.8%) | | | |
|---|---|---|---|---|---|---|---|---|---|---|---|---|
| | Alive (n[d]=66) | | Dead (n[d]=14) | | Alive (n[d]=24) | | Dead (n[d]=24) | | Alive (n[d]=23) | | Dead (n[d]=20) | |
| | n[e] | EFG[f] (%) | n[e] | EFG[f] (%) | n[e] | EFG[f] (%) | n[e] | EFG[f] (%) | n[e] | EFG[f] (%) | n[e] | EFG[f] (%) |
| FF/TT/WW/++/++/+I /CC/++ | 1 | 1.3 | | | | | | | | | | |
| FF/+T/WW/++/++/+I /CC/++ | 3 | 3.8 | | | | | | | | | | |
| FF/TT/WW/++/++/+I /+C/++ | 6 | 7.5 | | | | | | | | | | |
| FF/TT/WW/++/++/++/+C/++ | 7 | 8.8 | | | | | | | | | | |
| FF/+T/WW/++/++/+I /+C/++ | 12 | 15.0 | 1 | 1.3 | | | | | 1 | 3.1 | | |
| FF/+T/WW/++/++/++/+C/++ | 2 | 2.5 | | | | | | | | | | |
| FF/++/WW/++/++/+I /+C/++ | | | | | | | | | 5 | 15.5 | | |
| +F/+T/+W/++/+G/++/+C/++ | 2 | 2.5 | | | | | | | | | | |
| +F/+T/+W/++/+G/+I /+C/++ | 2 | 2.5 | | | | | | | | | | |
| FF/TT/WW/++/++/+I /++ /++ | 3 | 3.8 | 3 | 3.8 | | | | | | | | |
| FF/TT/WW/++/++/++/++/++ | 10 | 12.5 | 1 | 1.3 | 3 | 8.8 | 4 | 5.0 | 1 | 3.1 | 2 | 2.9 |
| FF/+T/WW/++/++/++/++/AA | | | | | 1 | 2.9 | | | | | | |
| FF/+T/WW/++/++/+I /++/+A | | | | | 2 | 5.8 | 1 | 1.3 | | | | |
| FF/+T/WW/++/++/II /++/++ | | | | | 3 | 8.8 | | | | | | |
| FF/+T/WW/++/++/+I /++/++ | 15 | 18.8 | 9 | 11.3 | 7 | 20.4 | 6 | 7.5 | 2 | 6.2 | 1 | 1.4 |
| FF/+T/WW/++/++/++/++/++ | 2 | 2.5 | | | | | 2 | 2.5 | 8 | 24.8 | 9 | 12.9 |
| FF/++/WW/++/++/+I /++/++ | 1 | 1.3 | | | | | | | 2 | 6.2 | 1 | 1.4 |

| | | | | | | | | | | | | |
|---|---|---|---|---|---|---|---|---|---|---|---|---|
| FF/++/WW/++/++/+I /++/+A | | | | | 3 | 8.8 | 3 | 3.8 | | | | |
| FF/++/WW/++/++/++/++/AA | | | | | | | 2 | 2.5 | | | | |
| FF/+T/WW/++/++/++/++/+A | | | | | 1 | 2.9 | 3 | 3.8 | | | | |
| FF/++/WW/++/++/++/++/++ | | | | | | | | | 4 | 12.4 | 4 | 5.8 |
| +F/+T/WW/++/++/+I /++/++ | | | | | | | 1 | 1.3 | | | | |
| +F/+T/+W/+P/+G/+I /++/++ | | | | | | | 2 | 2.5 | | | | |
| +F/+T/+W/+P/+G/++/++/++ | | | | | 4 | 11.7 | | | | | 1 | 1.4 |
| +F/++/+W/+P/+G/++/++/++ | | | | | | | | | | | 2 | 2.9 |
| Total | 66 | 82.5 | 14 | 17.5 | 24 | 70.0 | 24 | 30.0 | 23 | 71.2 | 20 | 28.8 |

[a] L213F, V410L, V1007G, I1011M, and T1520I were also genotyped but none of these amino acid substitutions was detected.

[b] G1: Adults of the first generation of the field-collected insects (larvae) were used for the bioassay.

[c] mortality: Mortality when insects were treated with 59 ng permethrin.

[d] n: number of mosquitoes tested for the genotyping.

[e] n: number of mosquitoes observed.

[f] EFG: Estimated Frequency of each Genotype in the population.

Table S4

Primers used for genotyping studies

| Target allele(s) | Primer name | Sequence | Utility | Approximate length of PCR product (bp) | Allele(s) determined |
|---|---|---|---|---|---|
| S66F | Ae61F6 | CGCCAATGTTTCCGTTCCAT | Forward | 208 | |
| | Ae61R3 | CTGATTCGCGTAATAGCTGTCG | Reverse | | |
| | Ae61R2 | TGTCGATATCCTCGAGAGGCGT | Sequencing | | S66F |
| L199F L213F | Ae213F1 | ATTCACCGGCATCTACACGTTC | Forward & Sequencing | 311 | L199F, L213F |
| | Ae213R9 | TCAACCCGACAAGTATTCCTAC | Reverse | | |
| A434T | Ae410F1[a] | TTACGATCAGCTGGACCGTG | Forward | 180 | |
| | Ae434R4[a] | TCAAAAGAATTCGCTCACCCG | Reverse | | |
| | Ae410F2[a] | ATCAGCTGGACCGTGGCA | Sequencing | | V410L, A434T |
| L982W, S989P V1016G | AaSCF20[b] | GACAATGTGGATCGCTTCCC | Forward | 589 | |
| | AaSCR21[b] | GCAATCTGGCTTGTTAACTTG | Reverse | | |
| | AaSCF9[b] | ACGGTGGAACTTCACCGACT | Sequencing | | L982W, S989P, A1007G, I1011M/V |
| | AaSCR22[b] | TTCACGAACTTGAGCGCGTTG | Sequencing | | V1016G |
| T1385I | Ae1385F1 | CTTCGTTGCTTCACTCTGTGGA | Forward | 107 | |
| | Ae1385R2 | TCATACCCTGCATACGGGACA | Reverse | | |
| | Ae1385R3 | ATACCCTGCATACGGGACATGG | Sequencing | | T1385I |
| F1534C T1539A | AaSCF7[b] | GAGAACTCGCCGATGAACTT | Forward | | |
| | AaSCR7[b] | GACGACGAAATCGAACAGGT | Reverse | | |

| | | | | | |
|---|---|---|---|---|---|
| | AaSCR8[b] | TAGCTTTCAGCGGCTTCTTC | Sequencing | | T1520I, F1534C, T1539A |
| V1703G D1763Y | AlSCF6 | TCGAGAAGTACTTCGTGTCG | Forward | 280 | |
| | AlSCR8 | AACAGCAGGATCATGCTCTG | Reverse | | |
| | Ae1703R1 | AGGATCATGCTCTGGCCGAA | Sequencing | | V1703G, D1763Y |

[a] These primers were used for the previous studies (2).

[b] These primers were used for the previous studies (1, 3).

Table S5

*Vssc* genes used for making phylogenic tree in Fig. 3*C*.

| Name | Haplotype | Genbank accession number | References |
|---|---|---|---|
| LVP | None | AAEL023266[a] | (2) |
| SMK | None | OM460722 | Current study |
| 1534C | F1534C | OM460730 | Current study |
| FC66 | S66F, F1534C | OM460727 | Current study |
| FC213 | L213F, F1534C | OM460726 | Current study |
| GY | V1016G, D1763Y | OM460723 | Current study |
| PG | S989P, V1016G | OM460731 | Current study |
| PGG | S989P, V1016G, V1703G | OM460732 | Current study |
| FTW | L199F, A434T, L982W | OM460729 | Current study |
| FWI | L199F, L982W, T1385I | OM460728 | Current study |
| PGC | S989P, V1016G, F1534C | OM460724 | Current study |
| FTWC | L199F, A434T, L982W, F1534C | OM460725 | Current study |
| Mex-03 | F1534C | OM460734 | (2) |
| Mex-06 | V410L, S723T, V1016I, F1534C | OM460735 | (2) |
| SP01 | S989P, V1016G, F1534C | OM460733 | (2) |
| MCNaeg-LIC | V410L, S723T, V1016I, F1534C | OM460736 | (2) |
| MCNaeg-C | V253F, M374I, G923S, F1534C | OM460737 | (2) |

[a] VectorBase accession number.

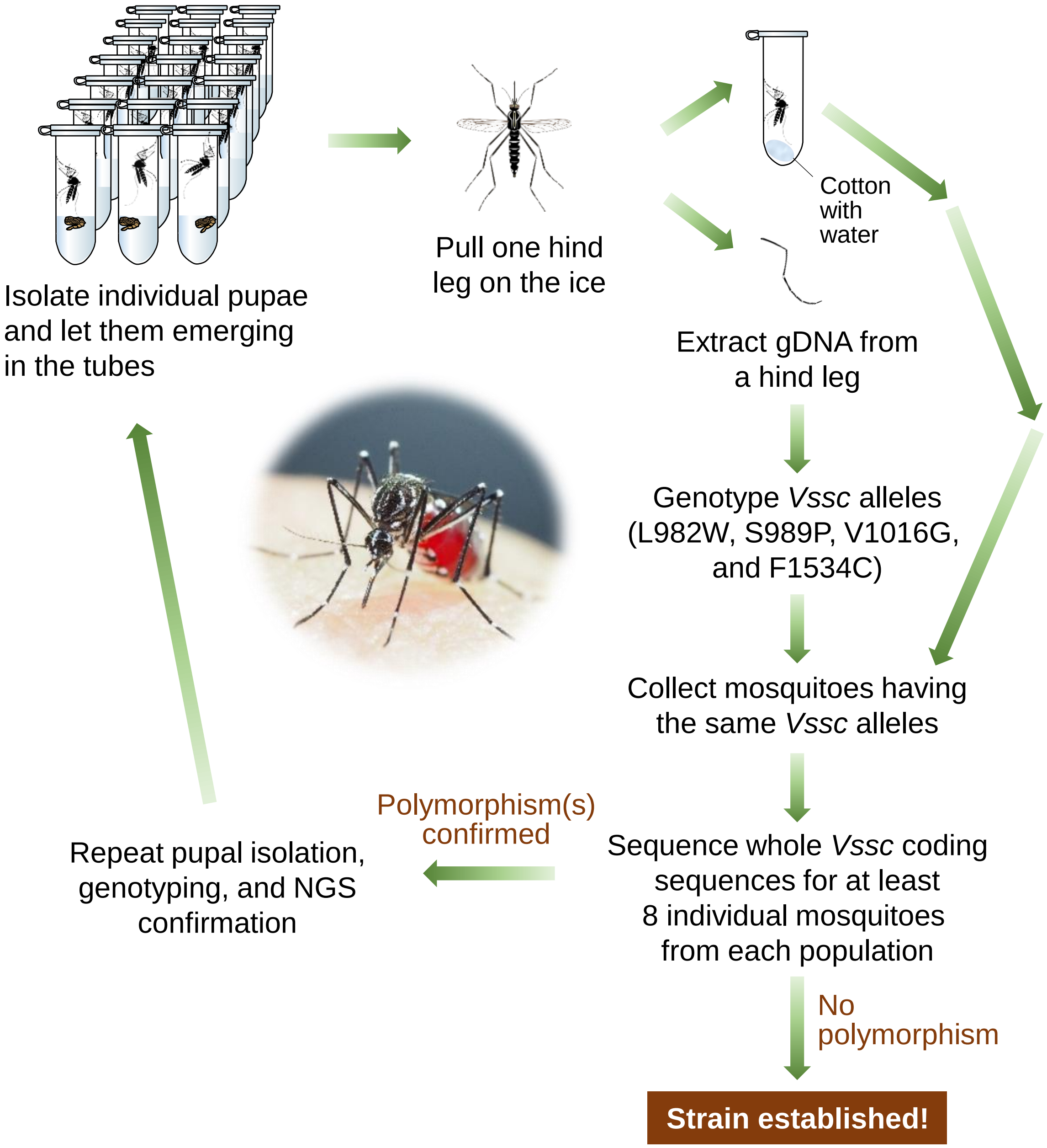


**Fig. S1.** Establishment of 10 knockdown resistance (*kdr*) strains of *Ae. aegypti*.

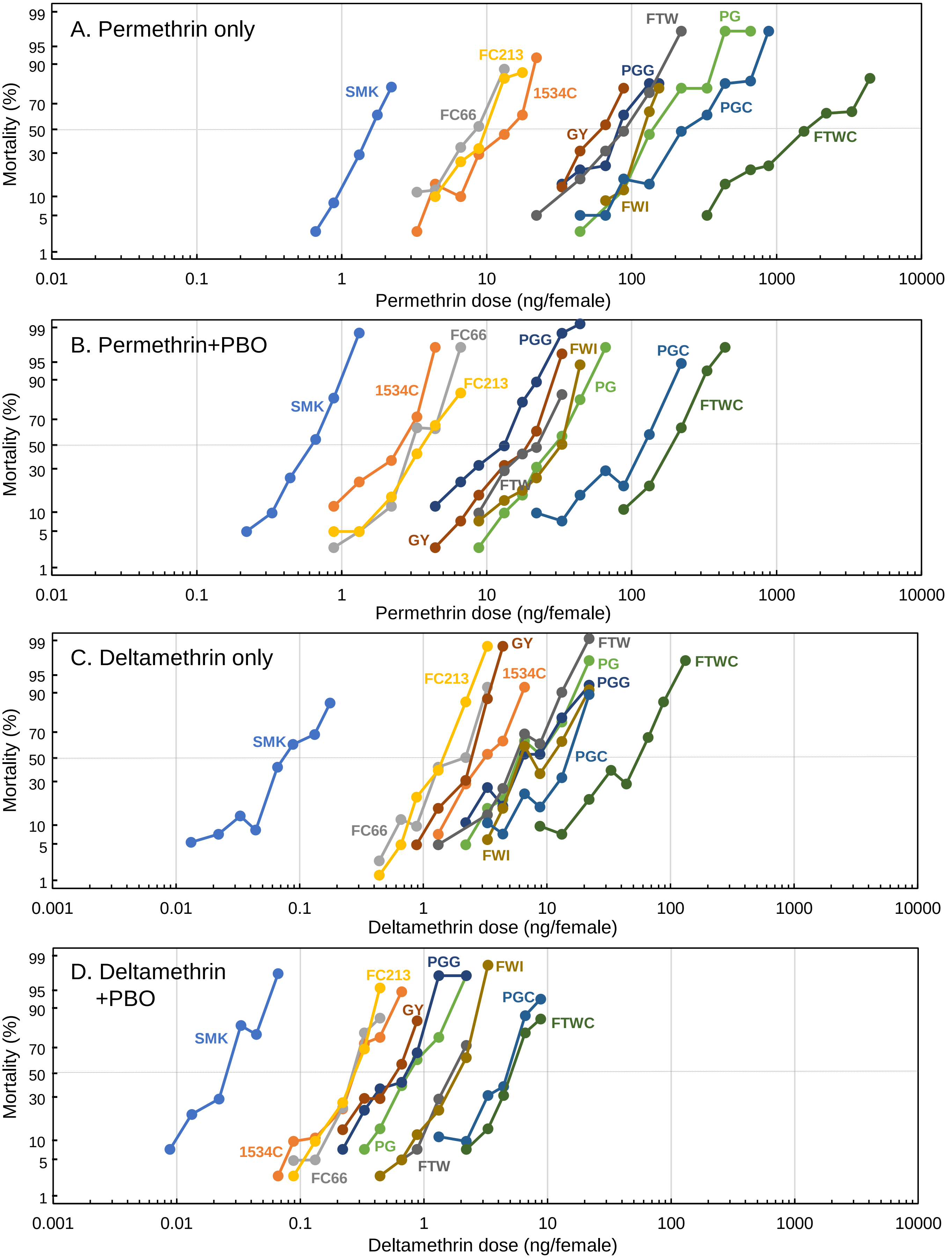


**Fig. S2.** Log dosage-probit mortality lines of eleven *Ae. aegypti* strains topically exposed to permethrin (A, B) and deltamethrin (C, D). Mosquitoes were treated with piperonyl butoxide (PBO) before applying pyrethroids to inhibit activity of cytochrome P450 monooxygenases (B, D).

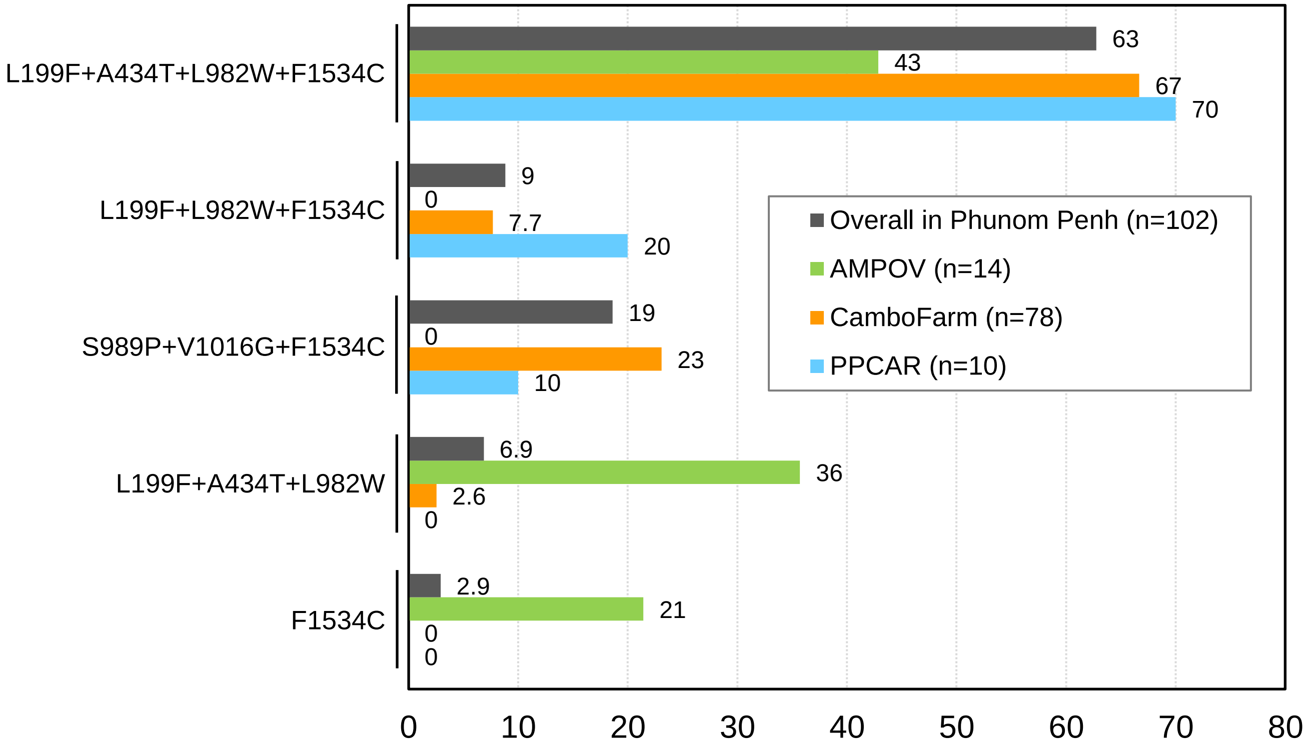


**Fig. S3.** Frequencies of 5 *Vssc* haplotypes in *Ae. aegypti* collected from Phnom Penh, Cambodia.

| Group | Species (accession) | L982**W**, inner ring, outer ring — DIIS6 |
|---|---|---|
| Insects | *Aedes aegypti* | DLPRWNFTDFMHSFMIVFRVLCGEWIESMWDCMLV-GDVSCIPFFLATVVIGN |
| | *Aedes albopictus* (AGE45998.1) | ***********************************-***************** |
| | *Acromyrmex charruanus* (KAG5347889.1) | ***********************************-***************** |
| | *Acromyrmex heyeri* (KAG5323878.1) | ***********************************-***************** |
| | *Acromyrmex insinuator* (KAG5314346.1) | ***********************************-***************** |
| | *Acromyrmex echinatior* (EGI61230.1) | ***********************************-***************** |
| | *Anopheles sinensis* (QGQ60479.1) | ***********************************-***************** |
| | *Anopheles gambiae* (CAA73920.1) | ***********************************-***************** |
| | *Anopheles arabiensis* (ADG29414.1) | ***********************************-***************** |
| | *Anopheles farauti* (ADM53043.1) | ***********************************-***************** |
| | *Anopheles stephensi* (ADZ96207.1) | ***********************************-***************** |
| | *Anopheles minimus* (ACX94090.3) | ***********************************-***************** |
| | *Anopheles funestus* (ABD59799.1) | ***********************************-***************** |
| | *Anopheles dirus* (AAY44402.1) | ***********************************-***************** |
| | *Apolygus lucorum* (ALF41049.1) | E********************************H*-***************** |
| | *Arctia plantaginis* (CAB3244532.1) | ***********************************-***************** |
| | *Aricia agestis* (XP_041980866.1) | ***********************************-***************** |
| | *Atta colombica* (XP_018045261.1) | ***********************************-***************** |
| | *Bactrocera oleae* (ABY21522.1) | *********************************Y*-***************** |
| | *Bicyclus anynana* (XP_023935270.1) | ***********************************-***************** |
| | *Bombyx mandarina* (XP_028027688.1) | ***********************************-***************** |
| | *Bombyx mori* (ACV87001.1) | ***********************************-***************** |
| | *Blattella germanica* (BBD13274.1) | *M*********************************-**W************** |
| | *Bradysia coprophila* (XP_037039280.1) | ***********************************-***************** |
| | *Brassicogethes aeneus* (AAO49190.1) | ***********************************-***************** |
| | *Brenthis ino* (CAH0724971.1) | ***********************************-***************** |
| | *Camponotus floridanus* (EFN61422.1) | EM*********************************-***************** |
| | *Cephus cinctus* (XP_024936215.1) | ***********************************-***************** |
| | *Contarinia nasturtii* (XP_031636803.1) | *M*********************************-***************** |
| | *Ctenocephalides felis* (QKK35480.1) | E**********************************-***************** |
| | *Culex erythrothorax* (QZU26835.1) | ***********************************-***************** |
| | *Culex pipiens* (ABM26922.1) | ***********************************-***************** |
| | *Culex pipiens pallens* (ADB13174.1) | ***********************************-***************** |
| | *Culex quinquefasciatus* (ABD64131.1) | ***********************************-***************** |
| | *Culex tarsalis* (QZU26834.1) | ***********************************-***************** |
| | *Culex tritaeniorhynchus* (AHV90608.1) | ***********************************-***************** |
| | *Cydia pomonella* (AFN44014.1) | ***********************************-***************** |
| | *Danaus chrysippus* (CAG9566212.1) | ***********************************-***************** |
| | *Drosophila albomicans* (XP_034119279.1) | *********************************Y*-***************** |
| | *Drosophila miranda* (XP_033242319.1) | *********************************Y*-***************** |
| | *Drosophila suzukii* (QBZ96182.1) | *********************************Y*-***************** |
| | *Eciton burchellii* (KAH0944544.1) | ***********************************-***************** |
| | *Fopius arisanus* (XP_011306710.1) | ***********************************-***************** |
| | *Formica exsecta* (XP_029662429.1) | E**********************************-***************** |
| | *Grapholita molesta* (ADD82833.1) | ***********************************-***************** |
| | *Habropoda laboriosa* (KOC69810.1) | ***********************************-***************** |
| | *Haematobia irritans* (AAC12793.1) | E********************************Y*-***************** |
| | *Helicoverpa armigera* (PZC85249.1) | ***********************************-***************** |
| | *Helicoverpa zea* (AXN77388.1) | ***********************************-***************** |
| | *Hermetia illucens* (XP_037914882.1) | ***********************************-***************** |
| | *Hyposmocoma kahamanoa* (XP_026326257.1) | ***********************************-***************** |
| | *Lucilia sericata* (XP_037809027.1) | E********************************Y*-***************** |
| | *Manduca sexta* (XP_037292327.1) | ***********************************-***************** |
| | *Melipona quadrifasciata* (KOX71756.1) | ***********************************-***************** |
| | *Monomorium pharaonic* (XP_012540554.1) | E**********************************-***************** |
| | *Musca domestica* (QCT80350.1) | E********************************Y*-***************** |
| | *Nasonia vitripennis* (XP_032452144.1) | E********************************R*-***************** |
| | *Nesidiocoris tenuis* (CAB0018697.1) | E********************************H*-***************** |
| | *Ochlerotatus trivitatus* (AAP60042.1) | ***********************************-***************** |
| | *Odontomachus brunneus (XP_032685106.1)* | ***********************************-***************** |
| | *Ooceraea biroi* (XP_026827710.1) | ***********************************-***************** |
| | *Osmia lignaria* (XP_034184267.1) | ***********************************-***************** |
| | *Ostrinia nubilalis* (ANW36302.1) | ***********************************-***************** |
| | *Papilio machaon* (XP_014357465.1) | ***********************************-***************** |
| | *Pararge aegeria* (XP_039758310.1) | ***********************************-***************** |
| | *Parnassius apollo* (CAG4965676.1) | ***********************************-***************** |
| | *Pieris macdunnoughi* (CAF4827565.1) | E**********************************-***************** |
| | *Plutella xylostella* (XP_037971760.1) | ***********************************-***************** |
| | *Pseudoatta Argentina* (KAG5322770.1) | ***********************************-***************** |
| | *Pulex irritans* (ANY39518.1) | E**********************************-***************** |
| | *Solenopsis invicta* (XP_039304537.1) | ***********************************-***************** |
| | *Spodoptera frugiperda* (AGK44162.1) | ***********************************-***************** |
| | *Spodoptera litura* (XP_022824852.1) | ***********************************-***************** |
| | *Stomoxys calcitrans* (XP_013100849.1) | E********************************Y*-***************** |
| | *Temnothorax longispinosus* (TGZ37383.1) | ***********************************-***************** |
| | *Tenebrio molitor* (KAH0816902.1) | E**********************************-***************** |
| | *Tuta absoluta* (AVG45379.1) | ***********************************-***************** |
| | *Vanessa tameamea* (XP_026493463.1) | ***********************************-***************** |
| | *Venturia canescens* (XP_043273604.1) | E********************************H*-***************** |
| | *Xenopsylla cheopis* (QLJ58300.1) | ***********************************-***************** |
| Mammals | *Mus musculus* (XP_006498646.1) | E****HMH**F***L******************E*A*QTM*LTV*MM*M**** |
| | *Rattus norvegicus* (EDM06395.1) | N****HMN**F***L****I*******T*****E*A*QAM*LTV*LMVM**** |
| | *Homo sapiens* (NP_001340889.1) | Q****HMN**F***L************T*****E*A*QAM*LTV*MMVM**** |

**Fig. S4.** Comparison of 81 amino acid sequences around outer and inner rings of the domain II from insects. Thirty one amino acids around the outer and inner rings are completely conserved among all species (boxed). Accession number of each protein is expressed in the parenthesis. Asterisks denote same amino acids as pyrethroid susceptible *Ae. aegypti*.

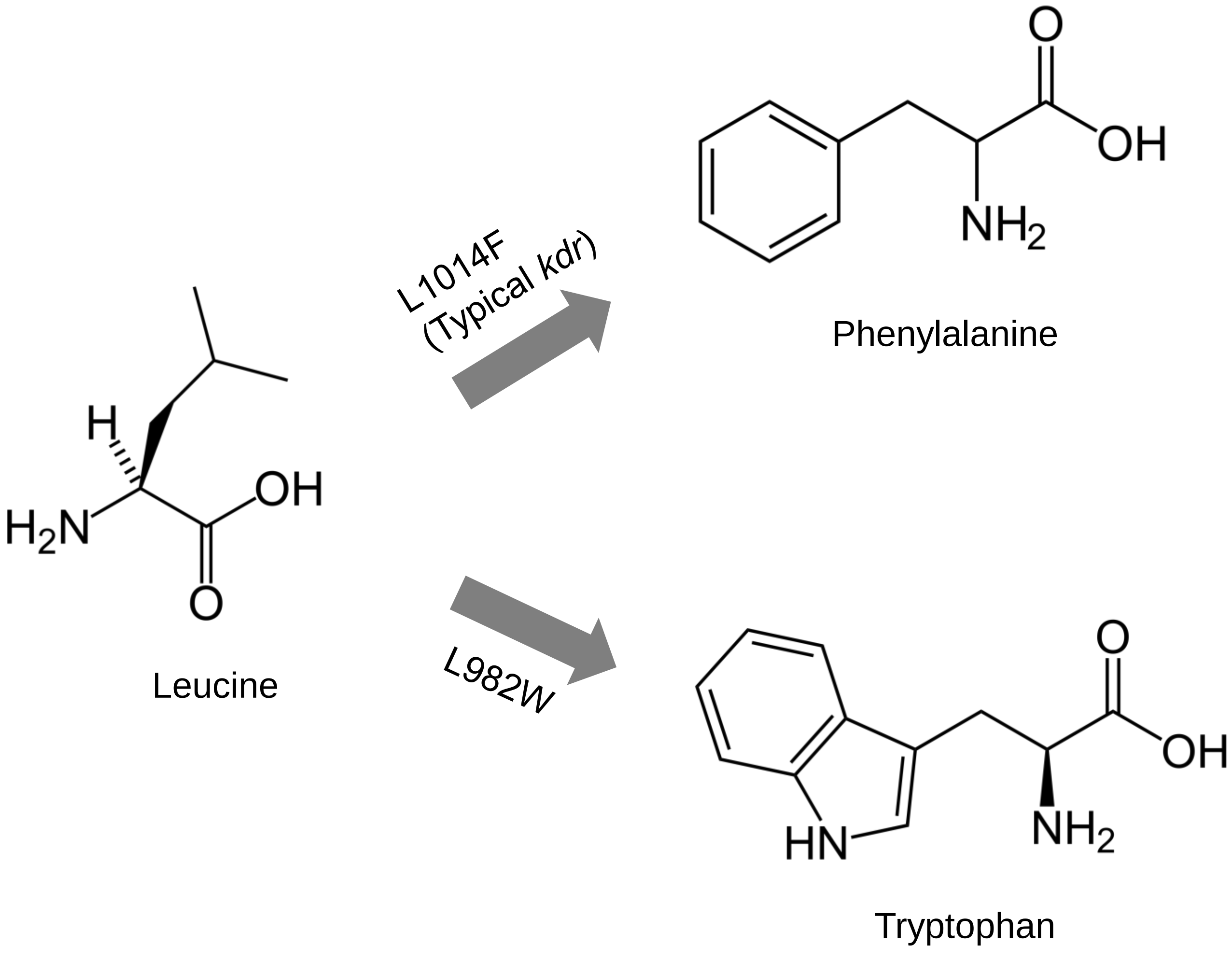


**Fig. S5.** Chemical structures of leucine, phenylalanine, and tryptophan.

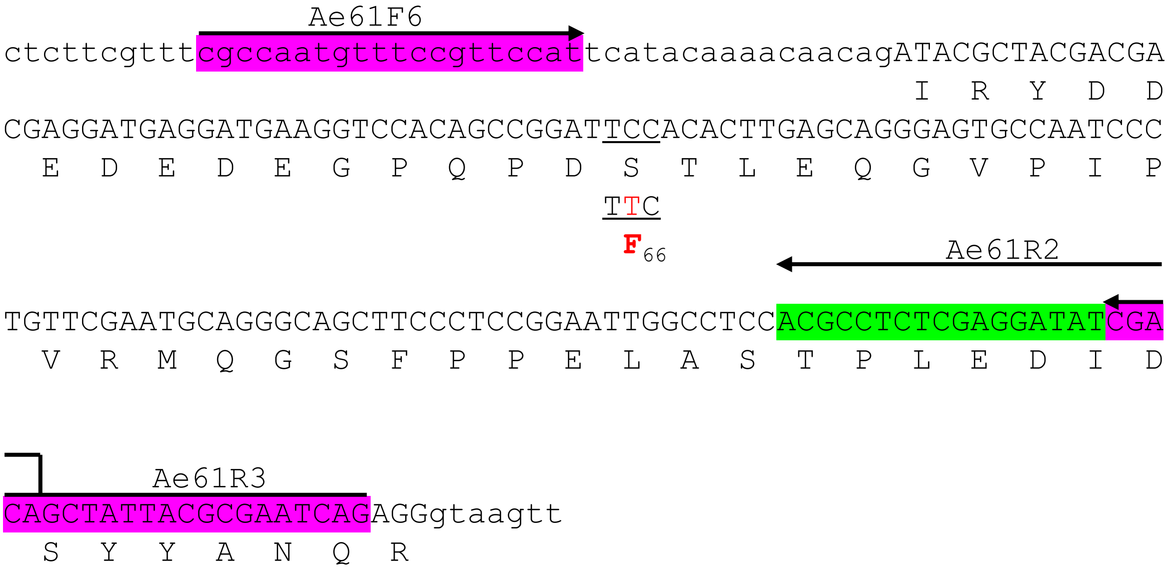


**Fig. S6.** Partial *Vssc* genome sequence of *Ae. aegypti* including S66F allele. Ae61F6 and Ae61R3 primers were used for PCR and Ae61R2 primer was used for sequencing. These primers were used to isolate FC66 and FC213 strains from HK-C (Fig. 2*B*). The S66F was not genotyped in the studies for Table 1 and Table S3 since it was unlikely that this allele conferred pyrethroid resistance.

```
                       Ae213F1
CCGAGGTGATATTCACCGGCATCTACACGTTCGAATCAGCTGTAAAAGTGATGGCGCGAG
   E  V  I  F  T  G  I  Y  T  F  E  S  A  V  K  V  M  A  R

GTTTCATATTACAACCGTTTACTTATCTTAGAGATGCATGGAATTGGTTGGACTTCGTAG
G  F  I  L  Q  P  F  T  Y  L  R  D  A  W  N  W  L  D  F  V
                          TTT
                           F199

TAATAGCATTAGCgtgagttttccagacactaatccttattgtgttcgatttttcttaat
V  I  A  L
        TTG
         F213

actccatatatacaatgctttaccccatataatcaactgtagtgtttaaagtgttatata

tagttcttatgttgattttgttttatttaaaacaagttttagttcattgtttgtaatctg

        Ae213R9
   *
taggaatacttgtcgggttgatatcagtcaa
```

*c in the LVP strain (VectorBase: AaegL5_3)

**Fig. S7.** Partial *Vssc* genome sequence of *Ae. aegypti* including L199F and L213F alleles. Ae213F1 and Ae213R9 primers were used for PCR and Ae213F1 primer was used for sequencing.

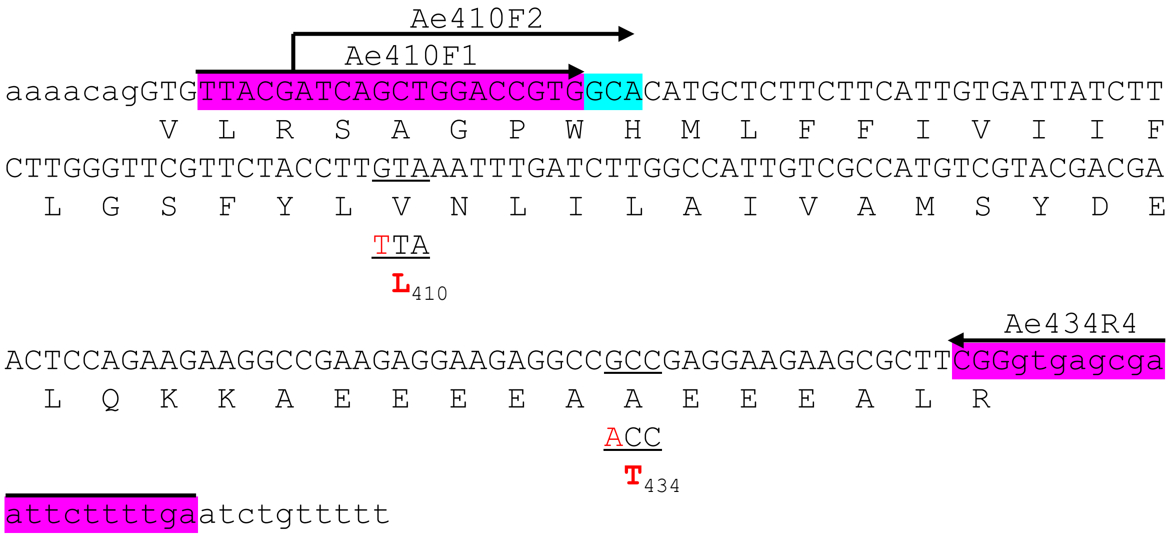


**Fig. S8.** Partial *Vssc* genome sequence of *Ae. aegypti* including V410L and A434T alleles. Ae410F1 and Ae434R4 primers were used for PCR and Ae410F2 primer was used for sequencing.

```
                AaSCF20 (→)                       AaSCF9 (→)
ctccccccagGACAATGTGGATCGCTTCCCGGACAAGGACCTGCCACGGTGGAACTTCACC
             N  V  D  R  F  P  D  K  D  L  P  R  W  N  F  T
GACTTTATGCACTCATTCATGATCGTGTTCCGGGTATTGTGCGGCGAGTGGATCGAATCC
 D  F  M  H  S  F  M  I  V  F  R  V  L  C  G  E  W  I  E  S
                                    TGG                  CCC
                                     W982                 P989
                                    CTG
                                     L982
ATGTGGGATTGTATGCTTGTGGGTGACGTGTCCTGTATTCCGTTCTTTTTGGCCACCGTA
 M  W  D  C  M  L  V  G  D  V  S  C  I  P  F  F  L  A  T  V
                                                   GGC
                                                    G1007
GTGATAGGAAATCTAGTAgtaagtattccgtttgggagttcatctataaggctgactgga
 V  I  G  N  L  V
   ATG
    M1011
   GTA
    V1011

agtaaattggagtgcacaacagacgtattatgctgtaattcgtgattcaactagttaaaa

tgaccgttgatcttgatagcatcaacactagaggcgtgctagcagcgagcgaggggcgta

ccaatttacttttggtcagcctttcttgcattctatcgtgctaaccgacaaattgtttcc

cacccgcacagGTACTTAACCTTTTCTTAGCCTTGCTTTTGTCCAATTTCGGTTCATCCT
            V  L  N  L  F  L  A  L  L  L  S  N  F  G  S  S
           GGA
            G1016
CGCTGTCGGCACCGACGGCCGACAACGAAACGAACAAGATCGCGGAGGCGTTCAATCGGA
S  L  S  A  P  T  A  D  N  E  T  N  K  I  A  E  A  F  N  R
                                            AaSCR22 (←)
TATCGCGCTTCTCCAACTGGATCAAGTCGAACATCGCCAACGCGCTCAAGTTCGTGAAAA
I  S  R  F  S  N  W  I  K  S  N  I  A  N  A  L  K  F  V  K
        AaSCR21 (←)
ACAAGTTAACAAGCCAGATTGCGTCCGTGCAG
N  K  L  T  S  Q  I  A  S  V  Q
```

**Fig. S9.** Partial *Vssc* genome sequence of *Ae. aegypti* including L982W, S989P, A1007G, I1011M/V, and V1016G alleles. AaSCF20 and AaSCR21 primers were used for PCR and AaSCF9 and AaSCR22 primers were used for sequencing. Non-synonymous polymorphisms were detected at the codon L982 (TTG and CTG).

```
                Ae1385F1
TCCTTAATCAACTTCGTTGCTTCACTCTGTGGAGCTGGTGGTATTCAAGCATTCAAAACA
 S  L  I  N  F  V  A  S  L  C  G  A  G  G  I  Q  A  F  K  T
                                                         ATA
                                                          I1385
                                                Ae1385R3
                                                   Ae1385R2
ATGCGAACTCTTAGAGCACTGAGACCGCTACGTGCCATGTCCCGTATGCAGGGTATGAGG
 M  R  T  L  R  A  L  R  P  L  R  A  M  S  R  M  Q  G  M  R

gtacgtag
```

**Fig. S10.** Partial *Vssc* genome sequence of *Ae. aegypti* including T1385I allele. Ae1385F1 and Ae1385R2 primers were used for PCR and Ae1385R3 primer was used for sequencing.

```
                 AaSCF7
CTACACGTGGGAGAACTCGCCGATGAACTTCGACCACGTGGGGAAGGCGTACCTGTGTCT
 Y  T  W  E  N  S  P  M  N  F  D  H  V  G  K  A  Y  L  C  L
GTTCCAGGTGGCAACGTTCAAGGGCTGGATCCAGATCATGAACGACGCCATCGACTCGCG
 F  Q  V  A  T  F  K  G  W  I  Q  I  M  N  D  A  I  D  S  R
GGAGgtaagttattgtgaaatcgaacttgttacgaatgatctgcttacaattttacgtcc
 E
tcgatccttccagGTGGGAAAGCAGCCGATTCGCGAGACCAACATCTACATGTACCTCTA
              V  G  K  Q  P  I  R  E  T  N  I  Y  M  Y  L  Y
                                     ATC
                                      I1520
CTTTGTGTTCTTCATCATCTTCGGGTCGTTCTTCACGCTGAATCTGTTCATCGGTGTCAT
 F  V  F  F  I  I  F  G  S  F  F  T  L  N  L  F  I  G  V  I
                   TGC            GCG
                    C1534          A1539
CATCGACAACTTCAACGAGCAGAAGAAGAAAGCCGGTGGCTCACTGGAAATGTTCATGAC
 I  D  N  F  N  E  Q  K  K  K  A  G  G  S  L  E  M  F  M  T

                                                  AaSCR8
GGAGGATCAGAAAAAGTACTACAACGCCATGAAAAAGATGGGCTCGAAGAAGCCGCTGAA
 E  D  Q  K  K  Y  Y  N  A  M  K  K  M  G  S  K  K  P  L  K

AGCTATTCCACGGCCTAGGgtaaggcatttccatcgcacatcaactgtgacgtattcctt
 A  I  P  R  P  R
cctaatctcgctattctcaatttcagTGGCGACCACAAGCAATAGTATTCGAAATAGTTA
                           W  R  P  Q  A  I  V  F  E  I  V
CCAATAAGAAGTTCGACATGATCATCATGTTGTTCATCGGGTTCAACATGTTGACGATGA
T  N  K  K  F  D  M  I  I  M  L  F  I  G  F  N  M  L  T  M
CGCTCGATCACTACAAGCAGACGGACACGTTTAGCGCGGTGCTAGACTATCTGAACATGA
T  L  D  H  Y  K  Q  T  D  T  F  S  A  V  L  D  Y  L  N  M
TCTTCATCTGCATCTTCAGTAGCGAGTGTCTGATGAAGATTTTCGCGCTGCGGTATCACT
I  F  I  C  I  F  S  S  E  C  L  M  K  I  F  A  L  R  Y  H
                          AaSCR7
ACTTTATCGAGCCGTGGAACCTGTTCGATTTCGTCGTCGTCATCCTGT
Y  F  I  E  P  W  N  L  F  D  F  V  V  V  I  L
```

**Fig. S11.** Partial *Vssc* genome sequence of *Ae. aegypti* including F1534C and T1539A alleles. AaSCF7 and AaSCR7 primers were used for PCR and AaSCR8 primer was used for sequencing.

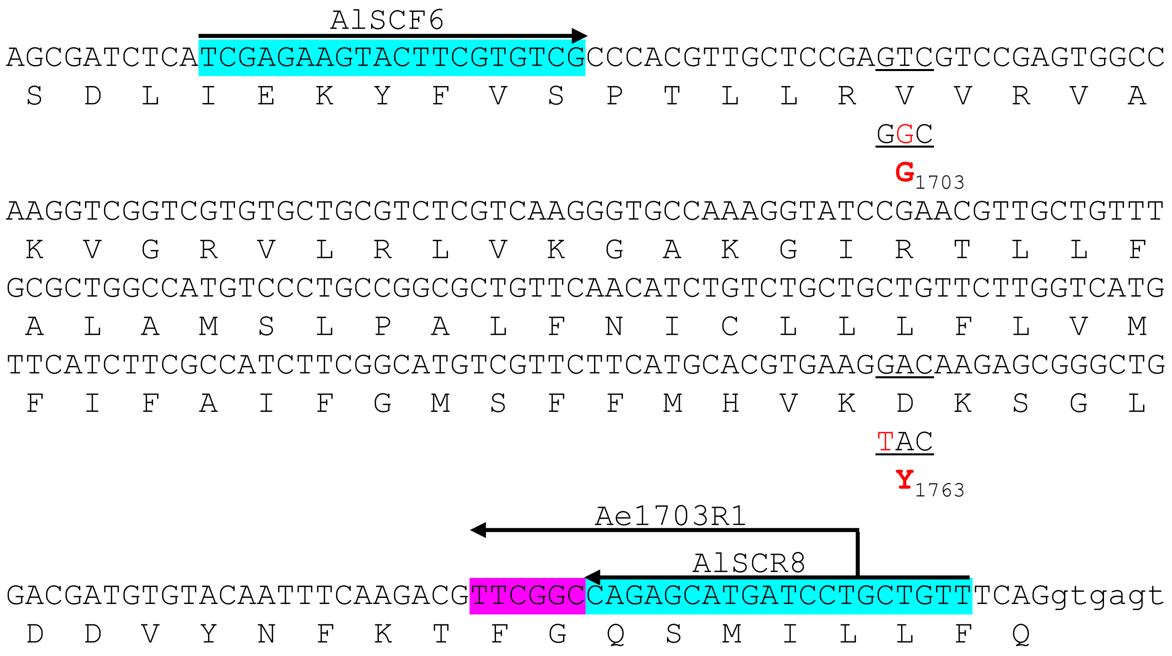


**Fig. S12.** Partial *Vssc* genome sequence of *Ae. aegypti* including V1703G and D1763Y alleles. AlSCF6 and AlSCR8 primers were used for PCR and Ae1703R1 primer was used for sequencing. These primers were used only for the preliminary experiments and V1703G and D1763Y were actually not genotyped in the studies for Table 1 and Table S3 since it was unlikely that these alleles conferred pyrethroid resistance.
